# Uncovering the Roles of *Mycobacterium tuberculosis melH* in Redox and Bioenergetic Homeostasis: Implications for Antitubercular Therapy

**DOI:** 10.1101/2023.10.02.560593

**Authors:** Yu-Ching Chen, Xinxin Yang, Nan Wang, Nicole S. Sampson

## Abstract

*Mycobacterium tuberculosis* (*Mtb*), the pathogenic bacterium that causes tuberculosis, has evolved sophisticated defense mechanisms to counteract the cytotoxicity of reactive oxygen species (ROS) generated within host macrophages during infection. The *melH* gene in *Mtb* and *Mycobacterium marinum* (*Mm*) plays a crucial role in defense mechanisms against ROS generated during infection. We demonstrate that *melH* encodes an epoxide hydrolase and contributes to ROS detoxification. Deletion of *melH* in *Mm* resulted in a mutant with increased sensitivity to oxidative stress, increased accumulation of aldehyde species, and decreased production of mycothiol and ergothioneine. This heightened vulnerability is attributed to the increased expression of *whiB3*, a universal stress sensor. The absence of *melH* also resulted in reduced intracellular levels of NAD^+^, NADH, and ATP. Bacterial growth was impaired, even in the absence of external stressors, and the impairment was carbon-source-dependent. Initial MelH substrate specificity studies demonstrate a preference for epoxides with a single aromatic substituent. Taken together, these results highlight the role of *melH* in mycobacterial bioenergetic metabolism and provide new insights into the complex interplay between redox homeostasis and generation of reactive aldehyde species in mycobacteria.

**Importance:** This study unveils the pivotal role played by the *melH* gene in *Mycobacterium tuberculosis* and *Mycobacterium marinum* in combatting the detrimental impact of oxidative conditions during infection. This investigation revealed notable alterations in the level of cytokinin-associated aldehyde, *para*-hydroxybenzaldehyde, as well as the redox buffer ergothioneine, upon deletion of *melH*. Moreover, changes in crucial cofactors responsible for electron transfer highlighted *melH*’s crucial function in maintaining a delicate equilibrium of redox and bioenergetic processes. MelH prefers epoxide small substrates with a phenyl substituted substrate. These findings collectively emphasize the potential of *melH* as an attractive target for the development of novel antitubercular therapies that sensitize mycobacteria to host stress, offering new avenues for combating tuberculosis.

---

*Mycobacterium tuberculosis* (*Mtb*) causes tuberculosis (TB), a significant global health threat that, in 2022, was the second leading cause of death from an infectious agent after COVID-19 (1). Within macrophages, the *Mtb* pathogen encounters formidable host cell defense responses, including nutrient limitation and redox stress. Despite these challenges, *Mtb* has developed mechanisms to evade macrophage antibacterial responses, adjusting its redox homeostasis and metabolic processes (2). The pathogen modulates its nutritional behavior and metabolic fluxes in response to different carbon sources during infection and growth (3). However, the precise role of *Mtb*’s metabolic flexibility in maintaining redox homeostasis remains unclear. Therefore, gaining a comprehensive understanding of the adaptive mechanisms employed by *Mtb* to survive in the human host holds the potential to significantly contribute to the identification of innovative therapeutic strategies.

In the *Mtb* genome, at least eight potential epoxide hydrolases (EHs) are identified, characterized by the presence of the αβ hydrolase domain (4). These EHs play a crucial role in converting epoxides to trans-dihydrodiols, believed to be essential for detoxification reactions necessary to withstand the hostile environment within host macrophages (5). Some EHs have been identified as potential therapeutic targets due to their involvement in detoxification processes (6–8). Previous evidence suggests that the *mel2* locus significantly contributes to oxidative stress resistance in *Mtb*-infected macrophages, although biochemical data regarding its function remain unclear (7). In this study, our focus was on characterizing epoxide hydrolase B (EphB or MelH) encoded by the *melH* (*Rv1938*) gene within the *mel2* locus. We successfully produced soluble MelH and demonstrated its epoxide hydrolase activity *in vitro* using a synthetic fluorescent substrate (PHOME) and screened additional epoxide substrate candidates to assess substrate specificity.

Several studies have unveiled *Mtb*’s sophisticated mechanisms for continuously monitoring and orchestrating appropriate responses against host-generated stresses. *Mtb* activates various transcriptional regulators in response to adverse conditions, exemplified by the control of various regulators to counteract oxidative stress. One such regulator is *whiB3*, a redox-sensing transcription factor encoding a 4Fe-4S redox sensor, that is sensitive to reactive oxygen species (ROS) (9). WhiB3 plays a pivotal role in maintaining intracellular redox homeostasis, ensuring both metabolic and cellular integrity (10). Mce3R, a TetR-type transcriptional repressor, controls the expression of *mel2* genes, including *melH* (11). A recent report identified an interrelationship between cholesterol, pH, and potassium levels dependent on Mce3R, emphasizing the crucial role of this regulon in host survival and its importance in responding to environmental stresses within the infected macrophage (12).

Previous studies have demonstrated that *whiB3* protects against the acidic pH encountered inside cells by modulating the mycothiol redox system (13). The *whiB3* repressor regulates the biosynthesis of both mycothiol (MSH) and ergothioneine (EGT) that serve as major redox buffers against various stressors (14) in mycobacteria. These redox systems contribute to mycobacterial survival strategies within the host (14).

In this study, we combined metabolomic, bioenergetic, biochemical, and transcriptomic approaches to evaluate how *melH* deletion impacts redox balance and bioenergetic homeostasis, which in turn moderates survival. Our investigation elucidated the critical importance of *melH* in regulating *whiB3*, maintaining MSH and EGT levels, and aldehyde accumulation, suggesting a link between all three in protecting against imbalanced redox stresses and in maintaining bioenergetic homeostasis.

## Results

### *melH-*encoded protein has epoxide hydrolase activity

*melH* (*Rv1938*) encodes a putative epoxide hydrolase B (EphB, MelH) with an alpha/beta hydrolase domain and a cap domain (8). Recombinant *Mtb* MelH was produced and purified (Figure S1A), and its identity was confirmed by mass spectrometry. We showed that a MelH hydrolyzed the artificial fluorogenic epoxide substrate PHOME (Figure S1B). The enzyme also demonstrated epoxide hydrolase activity similar to that of the C-terminal domains of a human soluble epoxide hydrolase (sEH) but lacked phosphatase activity (Figure S1C, D) (15).

### *melH* affects *Mtb* and *Mm* susceptibility to oxidative stress

Previous work has suggested that *mel2* locus is involved in the regulation of redox homeostasis (7, 16, 17). Through *in silico* analysis, it was discovered that among the six genes (melF-K) in the *mel2* locus of *Mm* and *Mtb*, *melF*, *melG*, and *melH* closely resemble homologs of bioluminescence-related genes, luxA, luxG, and luxH, respectively (16). The results of the study highlight the potential significance of *melF*, *melG*, and *melH* genes in the resistence of ROS-mediated oxidative stress. Here, we investigated how individual *mel2* gene mutations affected *Mtb*’s response to oxidative stress. Under normoxic conditions, *melF:Tn*, *melG:Tn*, and *melH:Tn* transposon mutants did not show increased ROS levels relative to the wild type (WT) in cultures grown with glycerol as sole carbon source. However, in response to potent oxidants, the *melH* mutant exhibited a significantly greater increase in ROS levels than the *melF* and *melG* mutants and wild-type (WT) *Mtb* CDC1551 (Figure 1A). The *melH* mutant also showed reduced survival rates compared to the WT and the *melF* and *melG* mutants under oxidative stress (Figure 1B). These findings suggest that *melH* plays a critical role in regulating ROS-mediated oxidative stress in *Mtb*.

**Figure 1.**
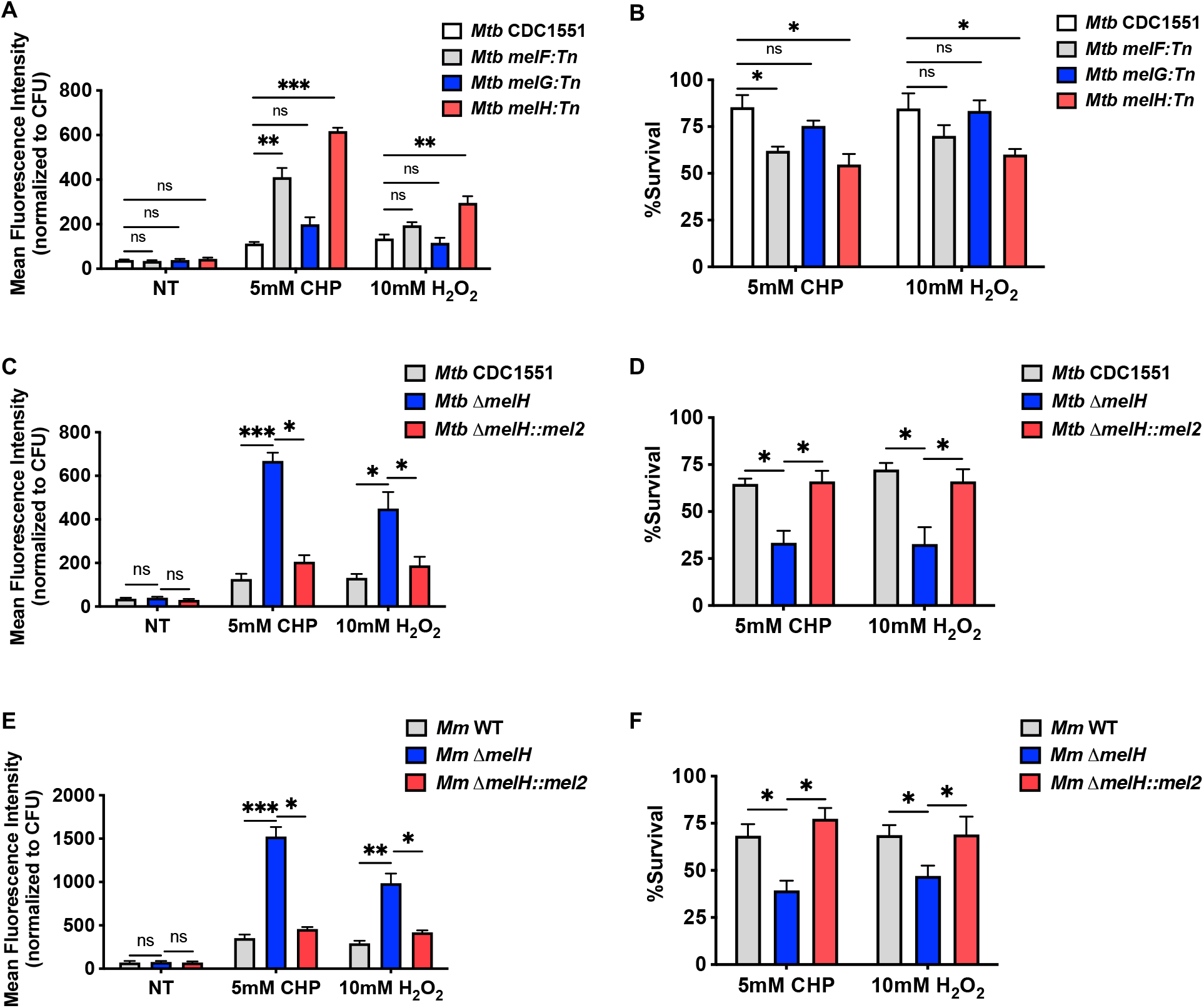
*melH* deletion exacerbates effect of and increases susceptibility to oxidative stress. (A) Mean fluorescence intensity of CellROX Green (an ROS-sensitive dye) in WT *Mtb* (CDC1551) and *melF/G/H* transposon mutants treated with CHP or H_2_O_2_. NT, no treatment. (B) Percentage survival of WT and mutant *Mtb* strains determined by measuring bacterial CFU counts after a 30-min treatment with CHP or H_2_O_2_ in Middlebrook 7H9 medium containing glycerol as a single carbon source. (C) Mean fluorescence intensity of CellROX Green in WT *Mtb* the Δ*melH* mutant, and the Δ*melH* complemented strain (*melH*::*mel2*) in 7H9 medium containing glycerol as a single carbon source. (D) Percentage survival of *Mtb* determined by measuring bacterial CFU counts after a 30-min treatment with CHP or H_2_O_2_ in 7H9 medium containing glycerol as a single carbon source. *P* values were determined by one-way analysis of variance using GraphPad Prism: \**P* < 0.05, \*\**P* < 0.01, \*\*\**P* < 0.001; ns, not significant. Error bars indicate standard deviations of three replicate experiments.

*Mycobacterium marinum* (*Mm*) has recently proven to be an applicable model for studying TB. Most importantly, the complete *mel2* locus, exhibiting gene arrangement and transcription orientation identical to the *Mtb*, has been identified in *Mm* (Figure S2A)(16). Using NCBI Protein BLAST (BLASTp), a sequence alignment was performed between the amino acid sequences of MelH from *Mtb* and *Mm*. The result showed that the two MelH sequences share a high degree of similarity (Figure S2B). Specificially, there was 87% amino acid identity, and 94% of the aligned positions showed positive matches, suggesting identity of physicochemical properties and function of MelH between *Mtb* and *Mm*.

To avoid any polar effects of transposition and to better understand the biochemical role of *melH* in *Mtb*’s defensive regulation of oxidative stress, we generated in frame Δ*melH* knock-out (KO) mutants and Δ*melH* complemented strains of *Mtb* and *Mm* by using specialized transduction in the clinical isolate *Mtb* CDC1551 and in *Mm* ymm1; and we validated the transformants by PCR (Figure S2C-F). First, we characterized the phenotypes of the Δ*melH Mtb* and *Mm* mutant and complemented strains (Figures 1C-F). Compared with ROS levels in WT *Mtb*, the levels in Δ*melH Mtb* were 3- to 4-fold and 2- to 3-fold higher in response to cumene hydroperoxide (CHP) and H_2_O_2_ treatment, respectively (Figure 1C and S3A). And the survival rate of Δ*melH* was lower than the rates of the WT and complemented strains upon treatment with CHP or H_2_O_2_ (Figure 1D and S3B). We observed an identical trend in *Mm* (Figure 1E and 1F). These results suggest that *Mtb* and *Mm* need *melH* to respond to ROS and to maintain redox balance, and utilize *melH* identically.

### Δ*melH* mutant exhibits altered growth profiles even in the absence of oxidative stress

*Mtb* utilizes metabolic pathways that include the tricarboxylic acid (TCA) cycle, pentose phosphate pathway (PPP), Embden–Meyerhof–Parnas (EMP) pathway, methyl citrate cycle, and the B12-dependent methylmalonyl pathway. These pathways equip *Mtb* to exploit a diverse array of carbon sources, including carbohydrates, sugars, fatty acids, amino acids, and sterols (18). While fatty acids from lipid droplets serve as the primary carbon source *in vivo*, host cells harbor various soluble nutrients that can function as alternative carbon sources (3). Our primary focus lies in comprehending how the *melH* mutant influences the modulation of metabolite flux within this network under distinct nutritional conditions. To understand the impact of different carbon sources on the metabolism of both WT *Mm* (ymm1), Δ*melH Mm*, and Δ*melH* complemented *Mm*, we cultured the mycobacteria in a carbon-defined minimal medium and examined their growth rates in the presence of diverse carbon sources. We observed altered growth phenotypes for both *Mm* anb *Mtb* Δ*melH* strains when they were grown with glycerol, propionate, cholesterol, or pyruvate as the sole carbon source (Figure 2A–C, F, and S4A-E). Interestingly, the growth rates of the two strains were similar when they were cultured with even-chained or long-chain fatty acids as the sole carbon source (Figure 2D, E, G and K). Also, the two strains exhibited similar growth rates when cultured with fumaric acid, succinic acid, or malic acid as the sole carbon source (Figure 2H-J). These results suggest that *Mm* and *Mtb* adapt their carbon flux in response to the environment, and the carbon-source-dependent growth-rate decrease observed in Δ*melH* emphasizes the importance of *melH* in central carbon metabolism. Given the identical phenotypes of *melH* in both *Mtb* and *Mm*, we performed subsequent experiments in *Mm* strains to facilitate experimental throughput.

**Figure 2.**
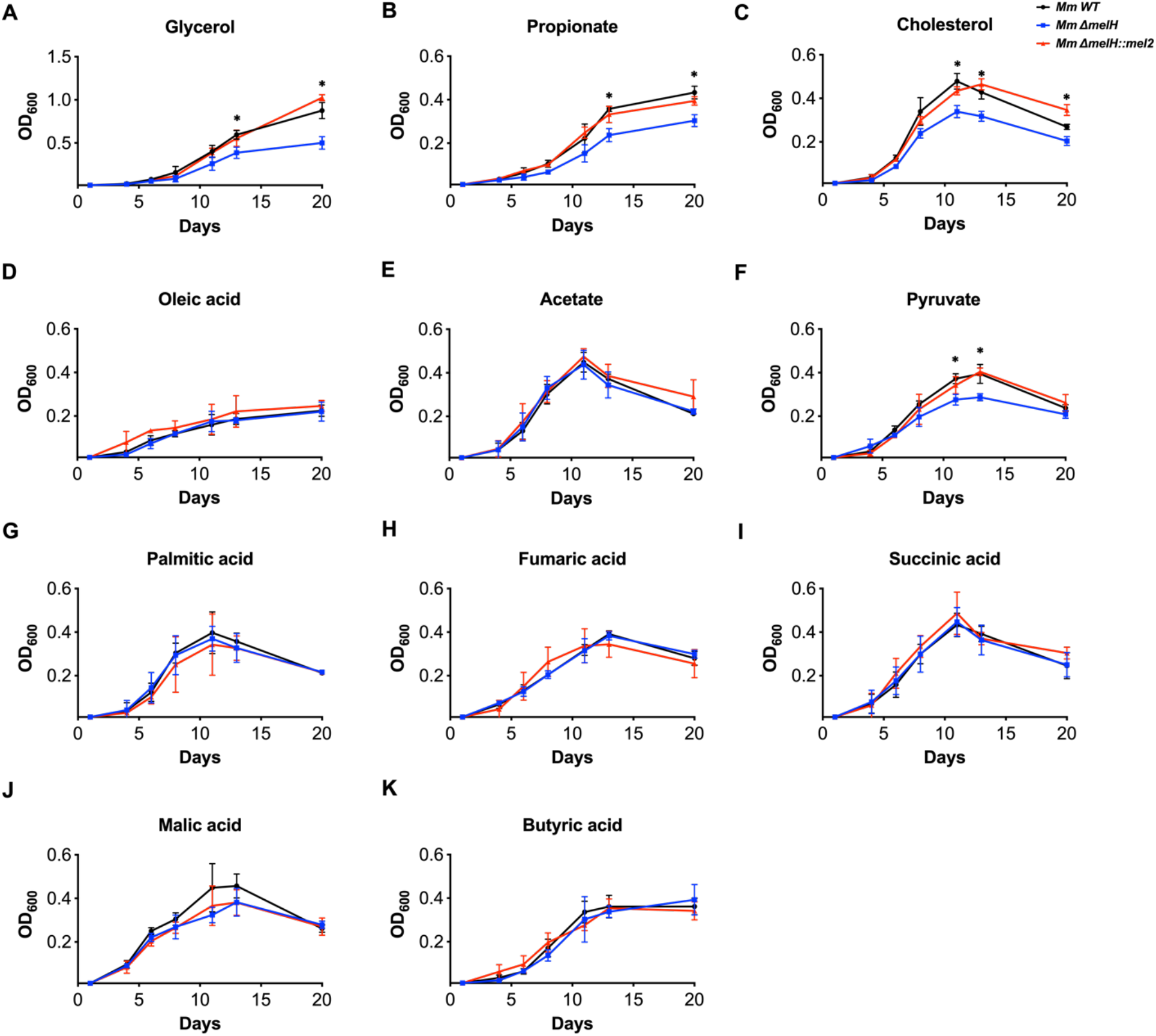
Effect of *melH* deletion on *Mm* growth rate depends on carbon source. Growth rates of WT *Mm* (black lines), the Δ*melH Mm* mutant (blue lines), and Δ*melH Mm* complemented (*melH*::*mel2*) (red lines) in minimal medium containing either (A) glycerol, (B) propionate, (C) cholesterol, (D) oleic acid, (E) acetate, (F) pyruvate, (G) palmitic acid, (H) fumaric acid, (I) succinic acid, (J) malic acid, or (K) butyric acid as a single carbon source. *P* values were determined by one-way analysis of variance using GraphPad Prism: \**P* < 0.05, \*\**P* < 0.01. Error bars indicate standard deviations of three replicate experiments.

### *melH* deletion modulates bioenergetic functions of *Mm*

In order to elucidate the molecular mechanisms underlying the carbon-source-dependent growth-rate defect and ROS-mediated oxidative damage observed in Δ*melH Mm*, we conducted untargeted metabolomics analyses of the WT, Δ*melH*, and Δ*melH* complemented strains grown in broth supplemented with glycerol, propionate, cholesterol, pyruvate, or oleic acid as the sole carbon source. The experimental design from the sample collection through data analysis is depicted in Figure S5A. The metabolites were extracted, separated by high-performance liquid chromatography, and subjected to electrospray ionization–time-of-flight mass spectrometry; and the metabolomes were analyzed with MetaboScape^®^. After evaluating heatmaps of differential abundance, we conducted a hierarchical clustering of m/z and retention time pairs, which revealed diverging sample clusters, with three groups mapped adjacent to each other (glycerol, propionate, cholesterol) and clearly separated from the oleic acid group (Figure S6A and S6B). Interestingly, in all groups of metabolomes, the differential abundance of metabolites between the WT and the mutant was minor. Of all the groups, the glycerol group showed the highest differential abundance between the WT and the mutant.

To gain a deeper understanding of the mechanisms linked to *melH* deficiency in *Mm*, we first selected the annotated and statistically significant metabolites observed in the tandem mass spectrometric confirmation experiments. Then we considered metabolites with an adjusted *P* value of <0.05 and a fold change of ≥|2.0| between WT and mutant for each carbon source to be differentially abundant metabolites (Figure 3A). The annotated and statistically significant metabolites were subjected to Metabolic Pathway Analysis (MetPA) (19) to assess changes in the abundance of metabolites from various biochemical pathways in Δ*melH Mm* compared to WT. The results showed the most statistically perturbed metabolic pathways to be nicotinamide metabolism, nucleotide metabolism, TCA cycle, and several amino acid biosynthesis pathways (Table 1 and Figure S7A).

**Figure 3.**
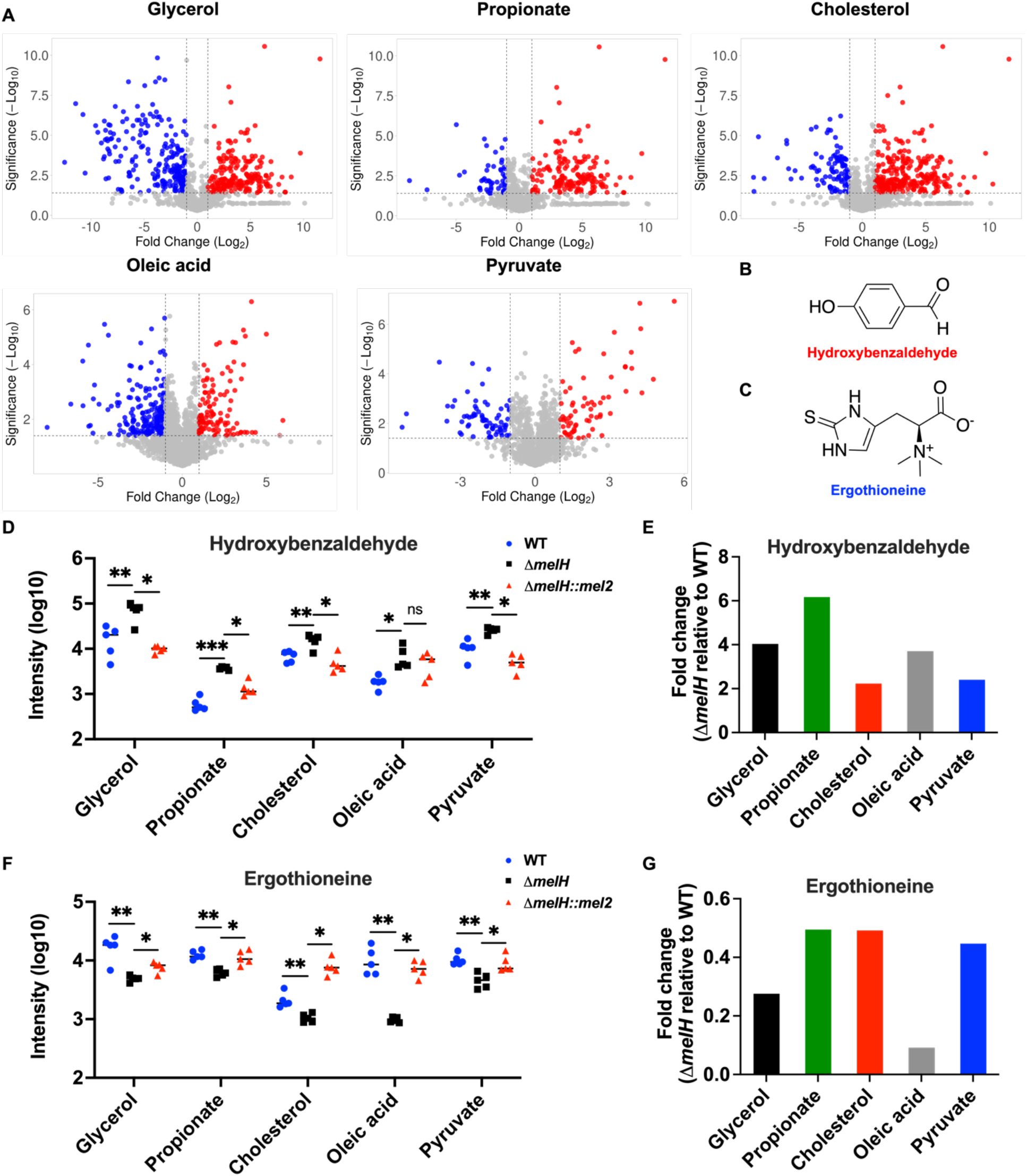
Liquid chromatography–mass spectrometry analysis of Δ*melH Mm* metabolites demonstrates increased *p*HBA levels and decreased EGT levels. (A) Volcano plots showing fold change versus significance for metabolites extracted from WT and Δ*melH Mm* strains cultured with glycerol, propionate, cholesterol, oleic acid, or pyruvate as the sole carbon source. Fold change of metabolite abundance in Δ*melH Mm* relative to the abundance in the WT. Red and blue dots indicate metabolites with ≥ 2.0-fold increase or decrease, respectively, on the *x*-axis and a corrected *P* value of <0.05 (-log^10^ > 1.3), on the y-axis; gray dots indicate metabolites with a <2.0-fold change and/or a *P* value of ≥ 0.05. (B and C) Chemical structures of (B) *p*HBA and (C) EGT. (D) Corresponding *p*HBA levels of in WT *Mm* (*N* = 5, blue dot), the Δ*melH Mm* mutant (*N* = 5, black square), and Δ*melH* complemented (*melH*::*mel2*) *Mm* (*N* = 5, red triangle). *P* values were determined by one-way analysis of variance using GraphPad Prism: \**P* < 0.05, \*\**P* < 0.01, \*\*\**P* < 0.001. (E) Fold change of EGT levels in Δ*melH Mm* relative to the level in the WT. (F) Corresponding EGT levels of in in WT *Mm* (*N* = 5, blue dot), the Δ*melH Mm* mutant (*N* = 5, black square), and Δ*melH* complemented (*melH*::*mel2*) *Mm* (*N* = 5, red triangle). *P* values were determined by one-way analysis of variance using GraphPad Prism: \**P* < 0.05, \*\**P* < 0.01, \*\*\**P* < 0.001. (G) Fold change of EGT levels in Δ*melH Mm* relative to the level in the WT.

**Table 1.**
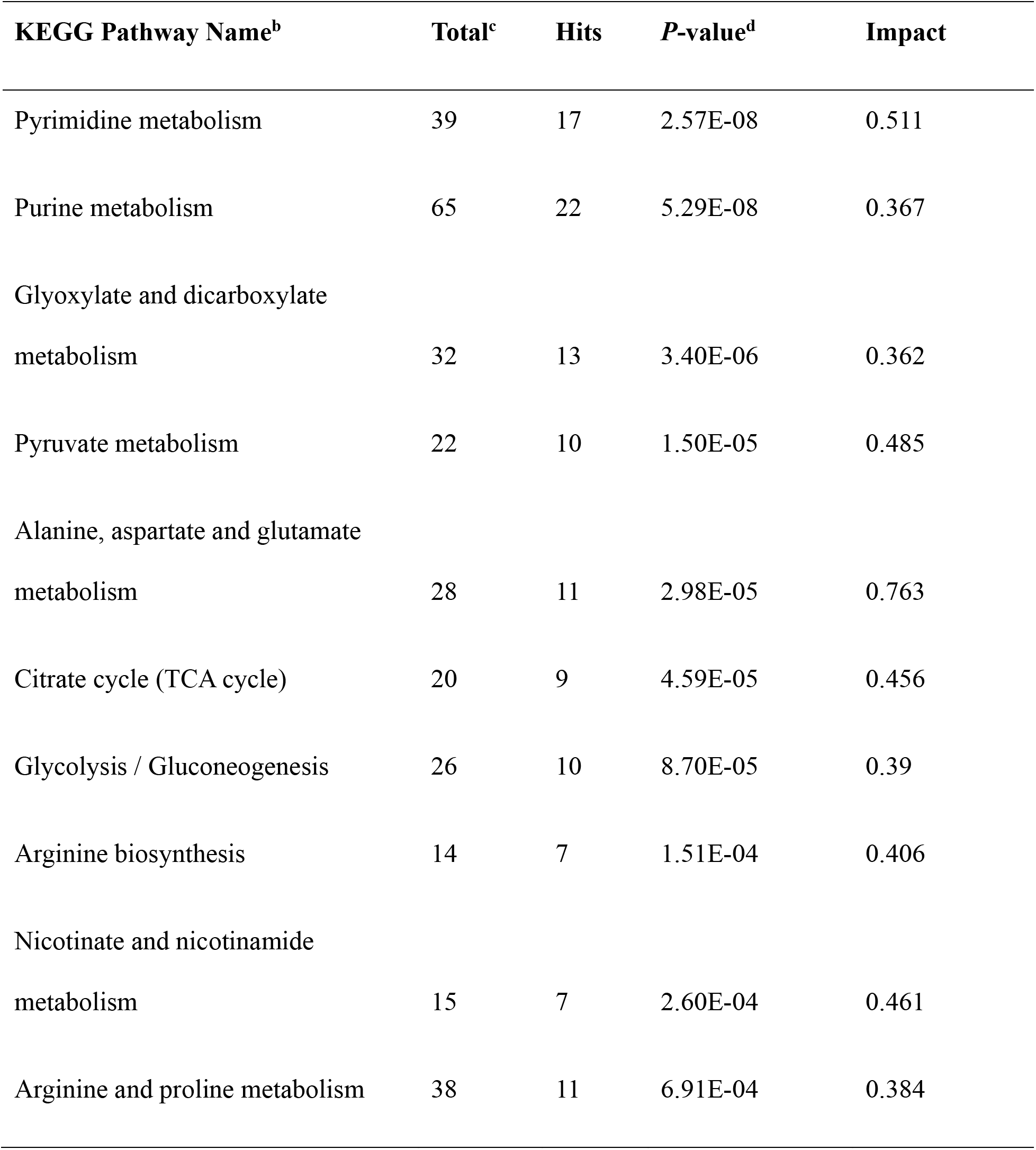
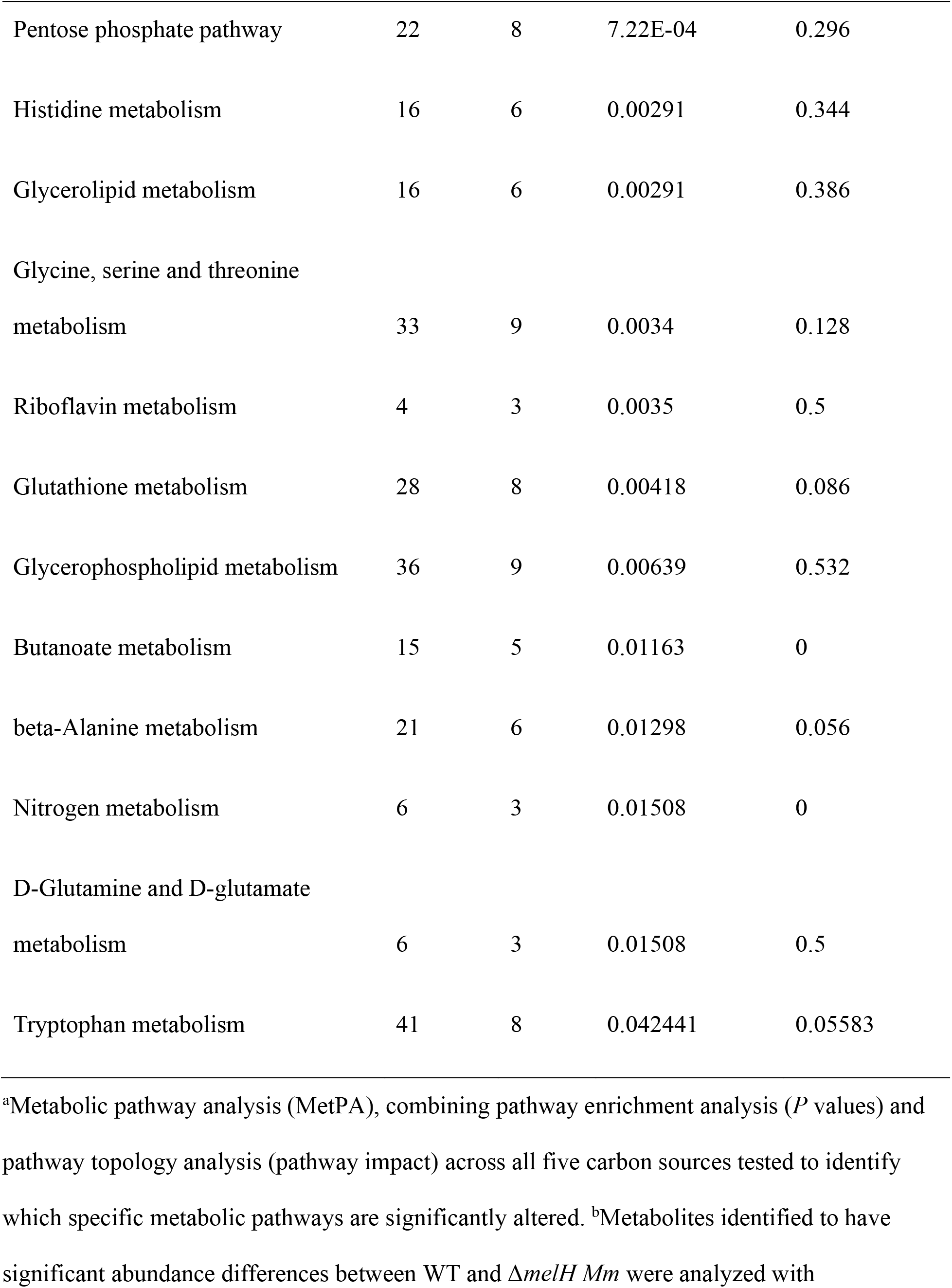
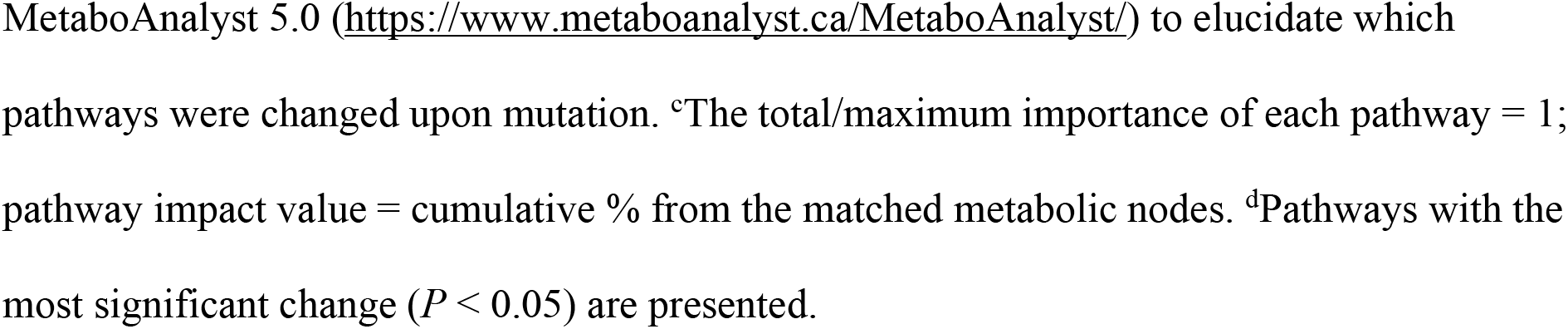
MetPa Analysis of metabolite pathways altered in △*melH* compared to WT *Mm*^a^.

We found lower abundance of NAD metabolite in the Δ*melH* mutant (Figure S7B). NAD acts as a key cofactor in many amino acid metabolism pathways, glycolysis, and the TCA cycle (20–23). Independent validation showed significantly lower intracellular concentrations of NAD+ and NADH (∼50%) in *ΔmelH Mm* compared to WT across most carbon sources (Figure 4A). When cholesterol was the exclusive carbon source a minimal difference between WT and mutant was exhibited in contrast to other carbon sources. Because the NAD^+^/NADH ratio serves as a critical indicator of cellular redox status, metabolic activity and cell health (24), we next asked whether *melH* deletion affected the NAD^+^/NADH ratio. Interestingly, the NAD^+^/NADH ratios in the WT and mutant did not differ significantly under any of the carbon source conditions, suggesting that *melH* deletion did not drive the cells into a more-reduced state in the absence of oxidative stress (Figure 4A).

**Figure 4.**
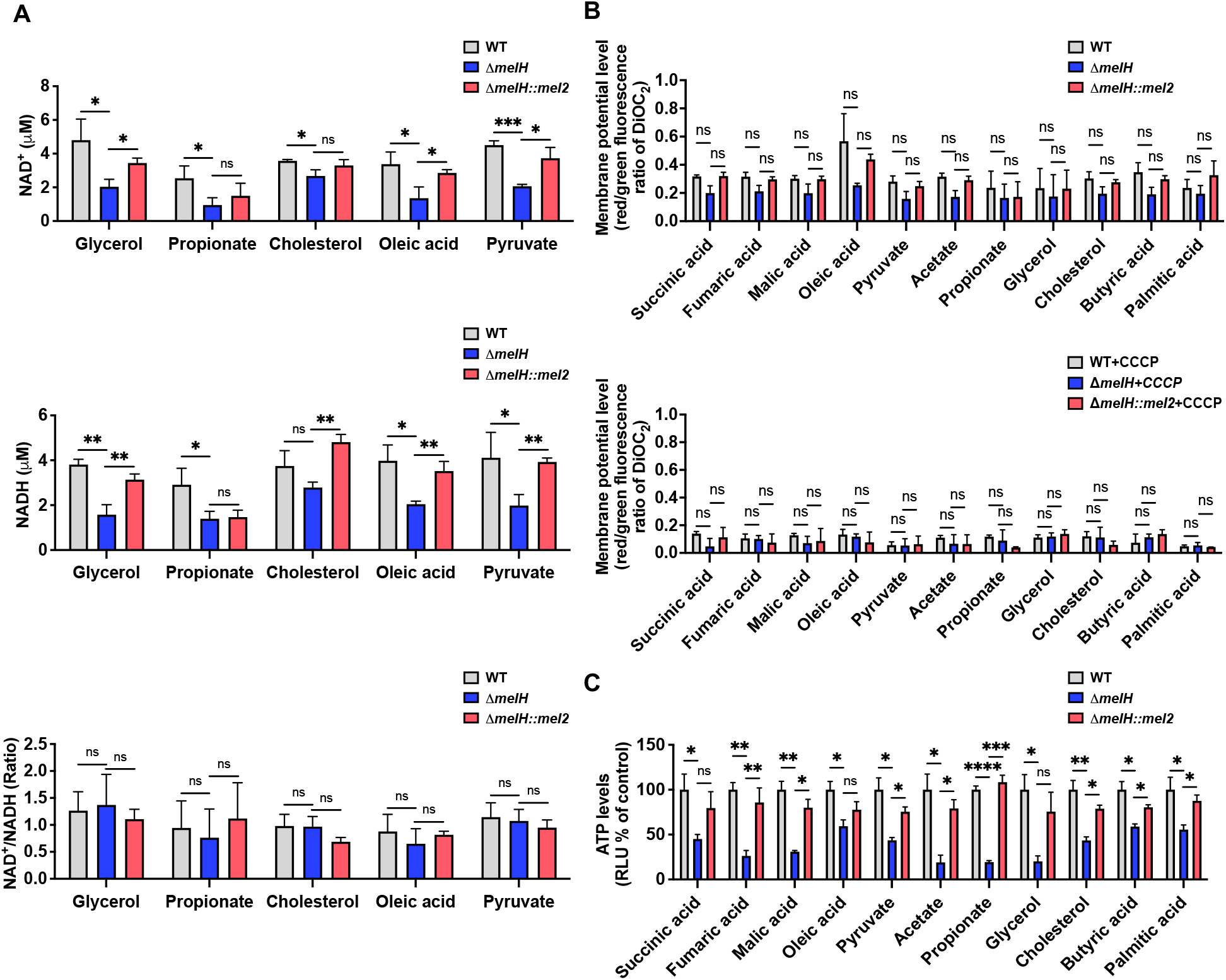
Deletion of *melH* reduces intracellular ATP levels and NAD^+^ and NADH concentrations in *Mm*. (A) Intracellular NAD^+^ and NADH concentrations in WT, Δ*melH*, and Δ*melH* complemented (*melH*::*mel2*) strains of *Mm*, as measured via recycling assays, along with calculated NAD^+^/NADH ratios. (B) Bacterial membrane potentials, measured with a BacLightTM Membrane Potential kit, for WT, Δ*melH*, and *melH*::*mel2 Mm* cultured with succinic acid, fumaric acid, malic acid, oleic acid, pyruvate, acetate, propionate, glycerol, cholesterol, butyric acid, or palmitic acid as the sole carbon source. DiOC_2_ = 3,3′-diethyloxacarbocyanine iodide. (C) Intracellular ATP levels, as measured by means of the luciferase/luciferin system, in WT, Δ*melH*, and *melH*::*mel2 Mm* cultured with succinic acid, fumaric acid, malic acid, oleic acid, pyruvate, acetate, propionate, glycerol, cholesterol, butyric acid, or palmitic acid as the sole carbon source. *P* values were determined by one-way analysis of variance using GraphPad Prism: \**P* < 0.05, \*\**P* < 0.01, \*\*\**P* < 0.001. Error bars indicate standard deviations of three replicate experiments.

Because electron transfer reactions are the main function of NAD (21), we next determined whether the lower intracellular NAD^+^ and NADH levels in the mutant might be due to increased proton conductance and resulting membrane depolarization, as seen with classical uncouplers. We used the fluorescent dye 3,3′-diethyloxacarbocyanine iodide (3) to measure the membrane potential of *Mm* cells. Δ*melH* showed no changes in membrane potential compared to WT under the tested carbon source conditions, although, as expected, the membrane uncoupler carbonyl cyanide *m*-chlorophenyl hydrazone (CCCP) did reduce the membrane potential compared to untreated mycobacteria (Figure 4B). Given this technique’s limit of detection for membrane changes, we cannot rule out a membrane potential change smaller than that caused by CCCP.

In mycobacteria, ATP synthesis depends primarily on the generation of proton motive force through the electron transport chain. Inhibition of the chain alters oxidative phosphorylation and ATP production (25). Since we have found lower metabolite abundance of ATP in the Δ*melH* mutant (Figure S7C). To directly address the bioenergetic status of the Δ*melH Mm* mutant, we measured intracellular ATP levels using a luciferase system. Independent validation showed that Δ*melH* had significantly lower intracellular ATP levels than WT (Figure 4C). Overall, the intracellular ATP level in the mutant was 3- to 4-fold lower than that in WT under most of the carbon source conditions. The difference in ATP level exceeded 4-fold when acetate, propionate, or glycerol was the sole carbon source. In contrast, a less than 2-fold difference was observed when oleic, butyric, or palmitic acid was the sole carbon source, consistent with lack of growth phenotype in even carbon-chain fatty acids. Although the NAD^+^/NADH ratio is not significantly altered, the reduced levels of NAD^+^ and NADH can still impact metabolic flux through redox reactions, potentially leading to decreased ATP levels. In sum, our results demonstrate that *melH* deletion lowered redox cofactor and energy storage levels in *Mm*.

### Metabolomic studies of Δ*melH Mm* show aldehyde accumulation and EGT depletion

Using the established cutoffs (Figure 3A), we further identified specific metabolites that were differentially abundant in the Δ*melH* mutant relative to WT across all tested carbon sources (Figures S6B and C). We observed increased levels of *para*-hydroxybenzaldehyde (*p*HBA) (Figure 3B, 3D and 3E), isonicotinic acid, and quinolinic acid (Figure S8C) and decreased levels of EGT (Figure 3C, 3F and 3G), L-acetylcarnitine, and tetrahydrofurfuryl butyrate (Figure S8D) in Δ*melH* relative to WT across all media.

The increase in *p*HBA observed in Δ*melH* compared to WT varies depending on the carbon source, with an approximately 2-fold increase in cholesterol or pyruvate medium, and approximately a 4-fold elevation in glycerol or oleic acid medium (Figure 3D and E). The highest increase was observed in propionate medium, with an approximately 6-fold increase in the mutant compared to WT. Additionaly, in Δ*melH*, we observed significant reductions in the levels of EGT in comparison to WT (Figure 3F and G). Specifically, EGT levels were decreased by approximately 10-fold when cholesterol was used as sole carbon source. When glycerol served as sole carbon source, EGT levels were reduced by about 3-fold. Similarly, in the presence of propionate, oleic acid, or pyruvate as sole carbon source, Δ*melH* showed a decrease in EGT levels, with an approximately 2-fold reduction in each case.

We also looked for MSH, a redox buffer distinct from EGT with a different reduction potential (26). We were only able to detect MSH in glycerol and pyruvate carbon sources (Figure S8D), it was absent in the remainder of our culture conditions. MSH is susceptible to facile oxidation under cell lysis conditions, whereas EGT is resistant to auto-oxidation, and predominantly exists in its thione form rather than the thiol form and has exceptional stability (27). These differences of EGT and MSH likely explain our inability to detect or quantify MSH metabolite in our metabolomics data.

### Aldehydes accumulate in *ΔmelH*

Darwin and coworkers have reported that adenine-based cytokinins and their degradation products, in particular *p*HBA, accumulate in an *Rv1205* mutant that sensitizes mycobacterial cells to NO (28–31). Given the accumulation of *p*HBA in Δ*melH Mm*, we examined the overall intracellular aldehyde levels in Δ*melH Mm*. Because aldehydes are inherently reactive and exist in different hydration states, their detection by mass spectrometry can be challenging. Therefore, we used a fluorgenic substrate that reacts with a broad range of aldehydes (32, 33). We measured intracellular aldehyde concentrations in *Mm* grown in glycerol, propionate, cholesterol, oleic acid, or pyruvate as the sole carbon source (Figure 5A). To ensure our assay was specific for aldehydes and not other reactive electrophiles, we tested glycidyl ether, an epoxide, as a control, and confirmed that epoxide does not react with the fluorogenic reagent.

**Figure 5.**
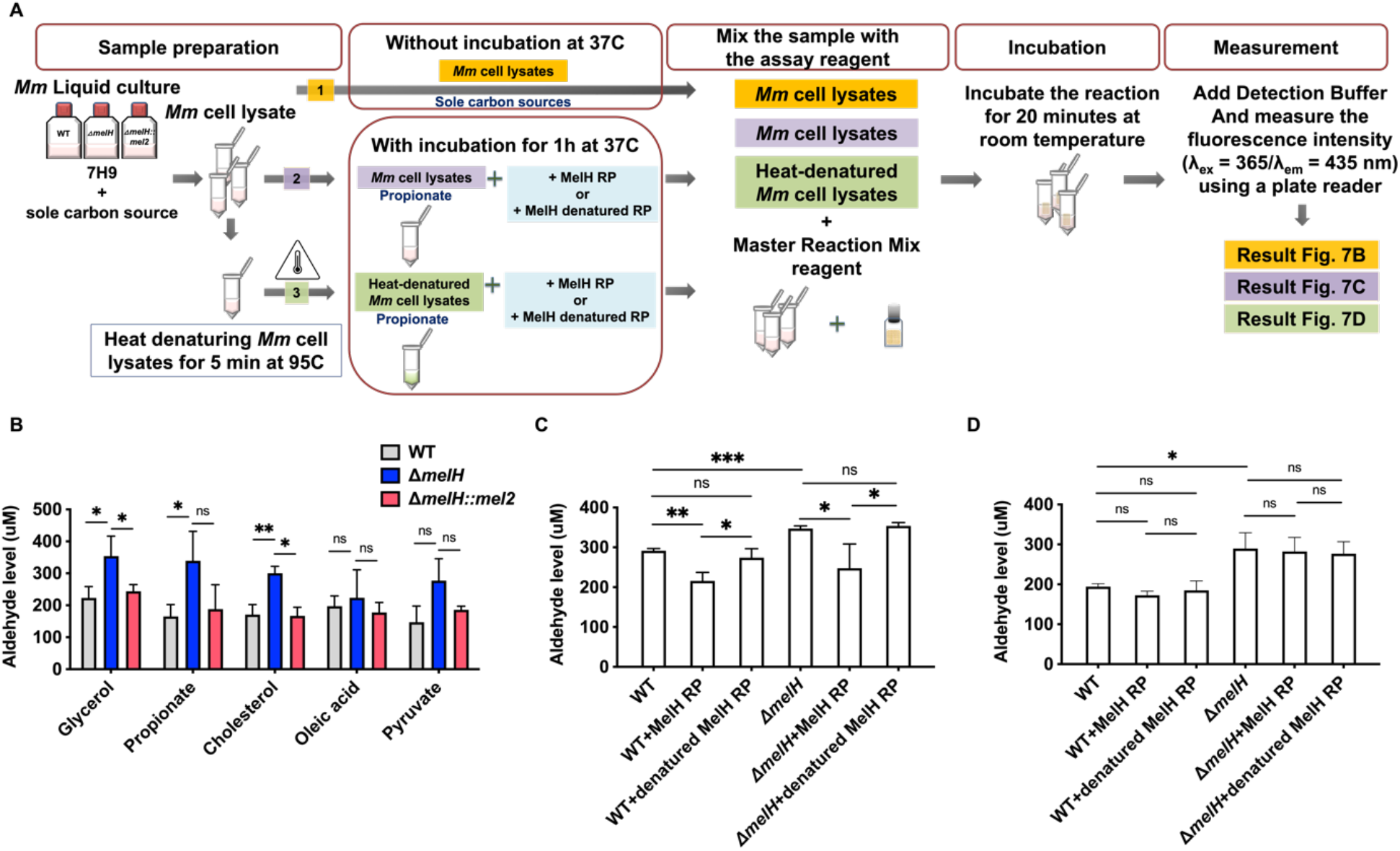
Disruption of *melH* causes intracellular aldehyde accumulation. (A) Workflow of aldehyde measurement. (B) Aldehyde concentrations in WT, Δ*melH*, and Δ*melH* complemented (*melH*::*mel2*) strains of *Mm* cultured with glycerol, propionate, cholesterol, oleic acid, or pyruvate as the sole carbon source. Aldehyde concentrations were determined with a fluorometric assay kit. After the mycobacteria were incubated with the aldehyde detection reagent for 30 min, the absorbance of the supernatants at 405 nm was measured using a plate reader. (C) Aldehyde concentrations in WT and Δ*melH Mm* lysates incubated with MelH recombinant protein or denatured MelH recombinant protein for 1 hour. (D) Aldehyde concentrations in enzyme-denatured WT and Δ*melH Mm* lysates incubated with MelH recombinant protein or denatured MelH recombinant protein for 1 hour. *P* values were determined by one-way analysis of variance using GraphPad Prism: \**P* < 0.05, \*\**P* < 0.01, \*\*\**P* < 0.001; ns, not significant. Error bars indicate mean ± SD (*N* = 3 independent experiments).

Δ*melH* showed significantly higher aldehyde concentrations when grown in glycerol, propionate, or cholesterol compared to the WT *Mm* and the Δ*melH* complemented strains. There were slightly higher, statistically insignificant concentrations of aldehyde in the mutant when grown in pyruvate; and no difference was observed when *Mm* was grown in oleic acid (Figure 5B). These results suggest that *melH* mutation alters aldehyde metabolism.

To explore further the role of MelH with respect to intracellular aldehyde concentrations, we determined whether exogenous recombinant MelH protein could prevent aldehyde generation in WT or Δ*melH Mm* cultured with propionate as the sole carbon source. We found that exogenous recombinant MelH reduced aldehyde concentrations in cell lysates of both strains, whereas heat-denatured recombinant MelH had no effect (Figure 5C).

In a subsequent experiment, we sought to investigate whether the accumulated aldehydes observed in the Δ*melH Mm* mutant were due to aldehyde-containing compounds serving as direct substrates for MelH or if MelH might function to eliminate an aldehyde precursor present in cell lysates. To address this question, we heat denatured cell lysates, and then added purified MelH protein to these cell lysates. If the accumulated aldehyde levels were solely due to the absence of catalytically-active MelH that formed aldehyde-containing compounds as its enzymatic product, then the addition of active MelH to the denatured lysate should have reduced the aldehyde concentrations. However, we observed that the aldehyde levels remained unchanged after the addition of MelH to denatured cell lysates (Figure 5D). This result strongly suggested that MelH’s direct substrate is not an aldehyde. Rather aldehydes are formed enzymatically from an unknown catalytic activity present in cell lysates and this enzymatic pathway utilizes as its substrate a metabolite that is depleted by addition of MelH epoxide hydrolyase catalyst to the cell lysate. This metabolite may be an epoxide or a precursor metabolite. In the absence of functional MelH in the Δ*melH Mm* mutant, these epoxides or precursors accumulate, and upon cell lysis, the aldehydes continue to be formed unless MelH is exogenously added to remove the epoxide/precursor. These results suggest a possible connection between MelH epoxide hydrolase activity and detoxification of intracellular aldehyde accumulation.

### Aldehyde accumulation minimally sensitizes *ΔmelH* to NO

In addition to enduring the presence of ROS, *Mtb* has developed the capability to resist host-generated antimicrobial molecules, including nitric oxide (NO). The precise mechanism of NO-mediated toxicity to *Mtb* remains unknown, but *Mtb* relies on a Pup-proteasome system (PPS) to withstand NO (34). Recent research has uncovered that pHBA, accumulates significantly in a PPS mutant. Notably, pHBA is sufficient to render *Mtb* sensitive to NO (35).

We next tested if aldehyde accumulation due to *melH* deletion correlated with increased susceptibility of the Δ*melH Mm* mutant to NO. *Mm* cells were exposed to acidified nitrite, a source of NO (36, 37). We observed that the nitrosative-stress-induced changes in nitrite concentration between the WT and the *ΔmelH* strain were slightly higher in the mutant, however, the differences were not statistically significant (Figure S9A). We also determined whether *melH* deletion reduced mycobacterial survival in *Mm* compared to WT upon induction of RNS. Acidified NaNO_2_ caused cell death at 30 min post-exposure as compared to unstressed bacteria; but the Δ*melH* survival rate was only slightly lower than the WT survival rate (Figure S9B). Complementation eliminated the slight reduction in Δ*melH* survival in response to acidified NaNO_2_. We also tested the effects of *S*-nitroso-N-acetylpenicillamine (SNAP) on the survival of the three strains and found that Δ*melH* was insignificantly more sensitive to SNAP than WT or the complemented strain. These results demonstrate that *melH* is not likely to be a major contributor to NO resistance in *Mm* despite the accumulation of pHBA.

### MelH prefers epoxide substrates with a single aromatic substitutent

We investigated the epoxide hydrolase activity of MelH on a set of potential epoxide substrates building on the previous work of James and coworkers (8). They found through activity assays and a liganded crystal structure that MelH has a preference for substrates with phenyl moieties. Consistent with their reports, UniProt bioinformatics similarity searches suggested the closest homologies were with enzymes that utilize aromatic substrates. Therefore, we selected the following five commercially-available substrates for our investigation: vitamin K1 2,3-epoxide, 2-biphenylyl glycidyl ether, styrene oxide, 1,4-naphthoquinone 2,3-epoxide, and (*E*)-1,3-diphenyl-2,3-epoxypropan-1-one. We tested each epoxide as an inhibitor and as a substrate of MelH. Inhibition was analyzed as percent reduction of turnover of the fluorescence substrate PHOME in the presense of a fixed concentration of epoxide. Substrate turnover was analyzed by analytical thin-layer chromatography (TLC) and ^1^H-NMR spectroscopy (Table 2). Styrene oxide (Figure S10) and vitamin K1 2,3-epoxide are completely converted to the corresponding diol. 1,4-Naphthoquinone 2,3-epoxide was only partially hydrolyzed after one hour of incubation with the enzyme, and the remaining two substrates were not hydrolyzed. Active substrates were poor inhibitors, presumably because they were rapidly converted to their corresponding product diols which do not bind as tightly to the enzyme. Interestingly, the three non-reactive or low-reactivity substrates, (*E*)-1,3-diphenyl-2,3-epoxypropan-1-one, 2-biphenylyl glycidyl ether and 1,4-naphthoquinone 2,3-epoxide inhibited MelH activity effectively (Table 2) suggesting that they bind non-productively to MelH due to their aromatic nature.

**Table 2.**
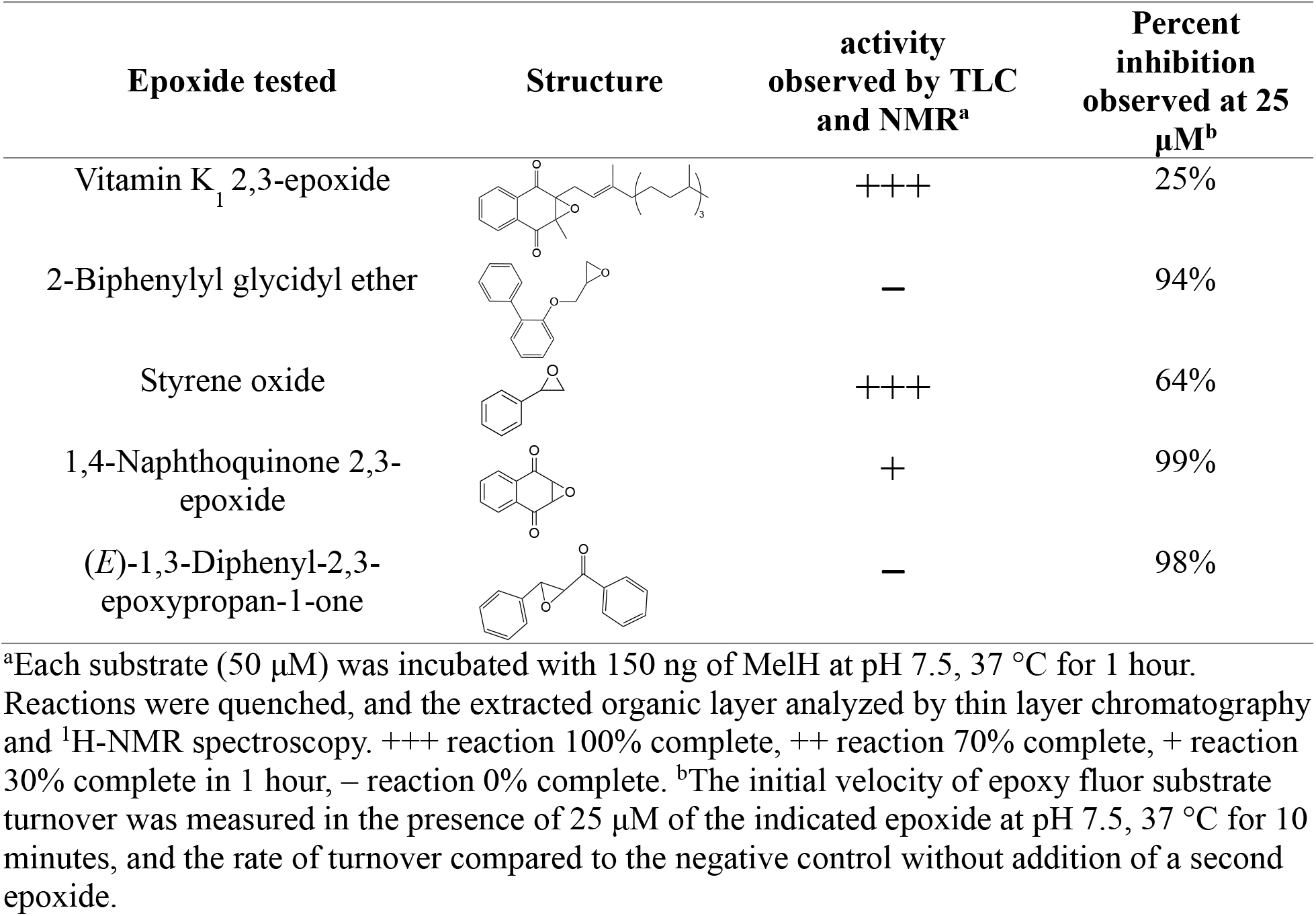
Evaluation of MelH potential epoxide substrates.

### *whiB3* regulates MSH and EGT production in response to oxidative stress

*Mtb whiB3*, an intracellular redox sensor, regulates EGT production in a carbon-source-dependent manner (14). EGT and MSH are the major redox buffer present in *Mtb* and Steyn and coworkers established that EGT and MSH are both required for maintaining redox balance and bioenergetic homeostasis, which influences drug susceptibility and pathogenicity (14). Our metabolomics analysis revealed that *melH* significantly decreased EGT and MSH levels (Figure 3F and S8D). Therefore, we compared the transcript levels for MSH and EGT biosynthesis genes MMAR_5212 (*egtD*), MMAR_5215 (*egtA*) and MMAR_0812 (*mshA*) as well as *whiB3 whiB3* (MMAR_1132) in the Δ*melH Mm* mutant relative to WT (38, 39) using quantitative real-time PCR.

Regardless of carbon source, deletion of *melH* resulted in an approximately two-fold upregulation of *whiB3* transcription compared with that in WT under normoxic conditions (Figure 6A). When treated with oxidative stressors, expression of *whiB3* was upregulated six to ten-fold in Δ*melH* compared with that in the WT in glycerol, propionate, cholesterol, pyruvate, or oleic acid culture conditions.

**Figure 6.**
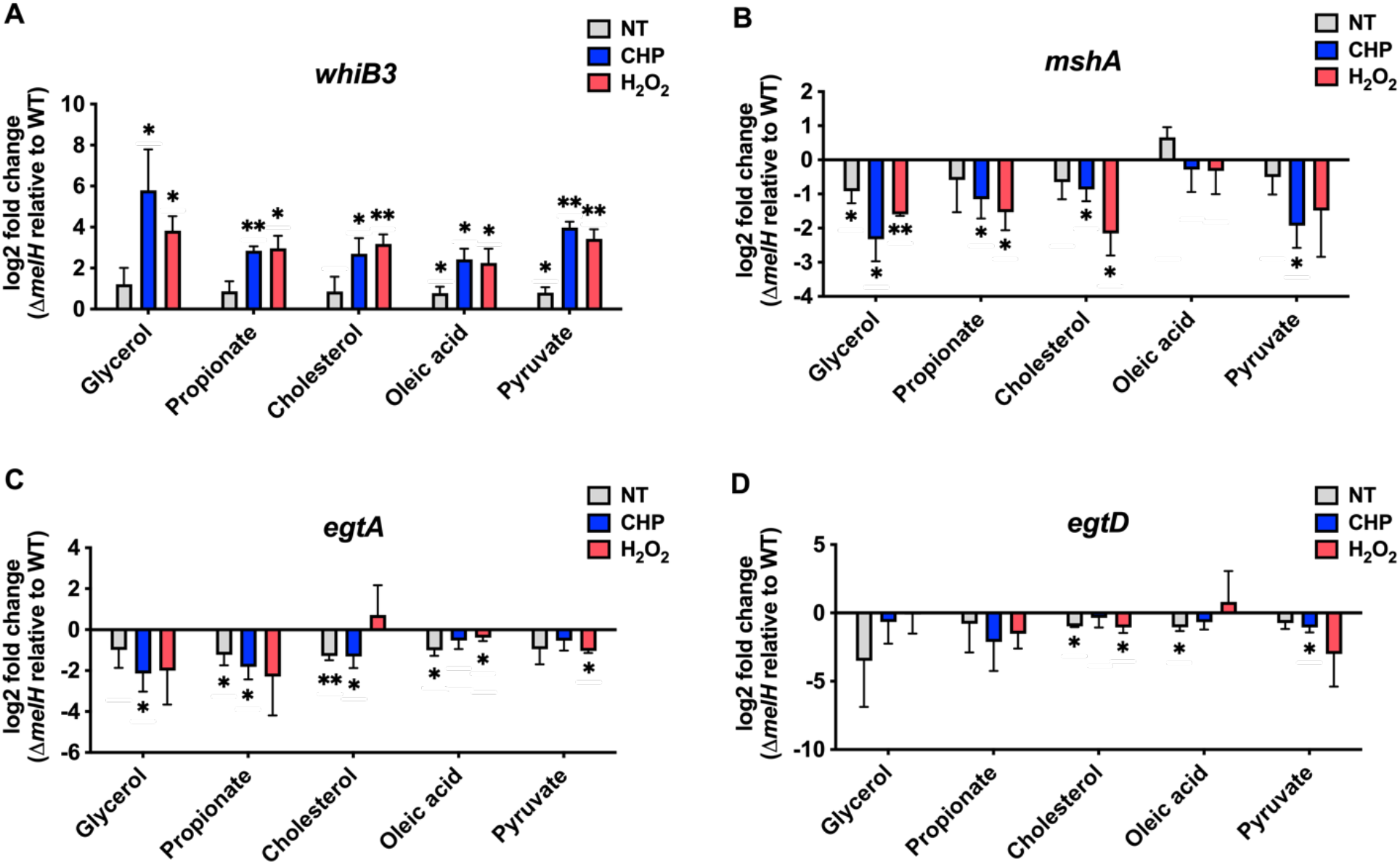
*melH* deletion affects expression of *whiB3* and EGT and MSH biosynthesis genes (*egtA*, *egtD*, and *mshA*) in *Mm* in response to oxidative stress. Comparison of (A) *whiB3*, (B) *mshA*, (C) *egtA*, and (D) *egtD* expression in *Mm melH* mutant relative to *Mm* WT under conditions of no treatment (NT), CHP, or H_2_O_2_. Total RNA was extracted from the bacteria, and intracellular gene expression was measured by quantitative real-time PCR. *P* values were determined by one-way analysis of variance using GraphPad Prism: \**P* < 0.05, \*\**P* < 0.01, \*\*\**P* < 0.001. Error bars indicate standard deviations of three replicate experiments.

In contrast, exposure of *Mm* to CHP or H_2_O_2_, significantly reduced expression of *mshA*, *egtA*, and *egtD* in the mutant compared to in WT (Figure 6B–D, respectively). Notably, when glycerol or propionate was the sole carbon source, the reduction in *egtA* expression in the CHP- or H_2_O_2_-treated mutant was more pronounced than the reduction observed under the other carbon source conditions. We observed significantly lower *mshA* expression in the Δ*melH* mutant than in the WT, particularly under oxidative stress conditions. Taken together, these results establish that *melH* deletion results in induction of *whiB3* expression and suggests that increased *whiB3* expression downregulates the expression of EGT and MSH biosynthesis genes.

### Δ*melH Mm* failure to maintain intracellular redox homeostasis is independent of carbon source

We examined the effect of *melH* deletion on redox homeostasis in *Mm* grown with different carbon sources and found that *melH* deletion resulted in increased ROS accumulation in response to oxidative stress regardless of carbon source. Specifically, Δ*melH* exhibited an approximately 1.5- to 3-fold increase in ROS production in response to CHP or H_2_O_2_ treatment across all carbon sources. Interestingly, the results do not show a significant change in the ROS production in Δ*melH* compared to the WT and the complemented strains under non-stressing growth conditions (Figure 7A).

**Figure 7.**
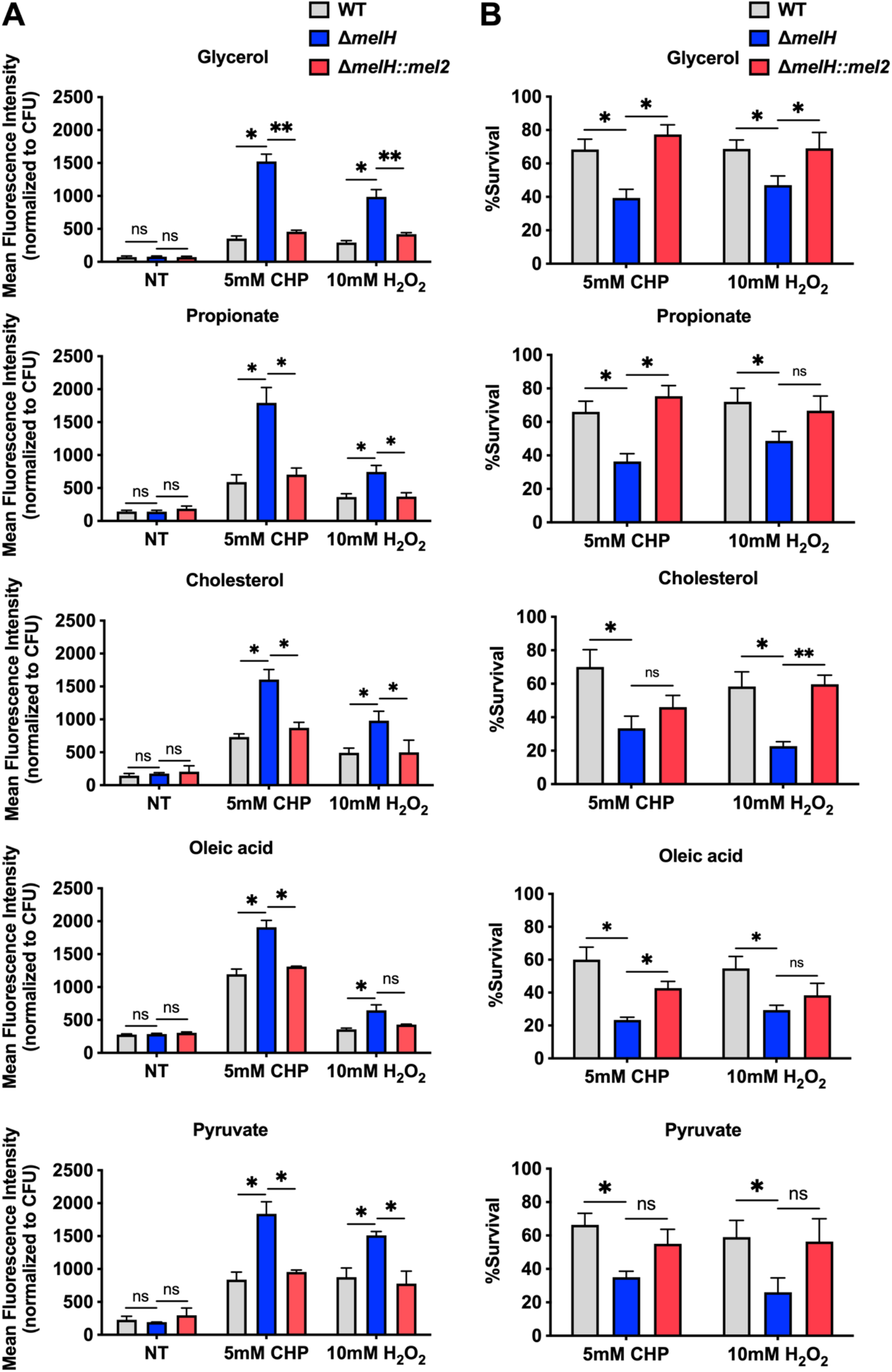
Absence of *melH* in *Mm* results in heightened vulnerability to oxidative stress in response to carbon catabolism. (A) Mean fluorescence intensity of CellROX Green (an ROS-sensitive dye) CHP- or H_2_O_2_-treated WT, Δ*melH*, and Δ*melH* complemented (*melH*::*mel2*) strains of *Mm* cultured in 7H9 medium with glycerol, propionate, cholesterol, oleic acid, or pyruvate as the sole carbon source. (B) Percentage survival of WT, Δ*melH*, and *melH*::*mel2* strains of *Mm* determined by measuring bacterial CFU counts after a 30-min treatment with CHP or H_2_O_2_ in 7H9 medium containing glycerol, propionate, cholesterol, oleic acid, or pyruvate as a single carbon source. *P* values were determined by one-way analysis of variance using GraphPad Prism: \**P* < 0.05, \*\**P* < 0.01, \*\*\**P* < 0.001; ns, not significant. Error bars indicate standard deviations of three replicate experiments.

To determine the impact of *melH*-mediated ROS toxicity on cellular survival, we measured CFUs after treatment of *Mm* cells with CHP or H_2_O_2_. We found that the survival rate of Δ*melH Mm* was significantly lower than the WT survival rate and that complementation of the mutant with the *mel2* operon restored intracellular survival to near WT levels across all carbon source growth conditions (Figure 7B). Altogether, the high expression of *whiB3* exhibited under CHP- or H_2_O_2_-induced oxidative stress conditions in Δ*melH* is consistent with the increased ROS levels and the decreased survival of the Δ*melH* mutant. The heightened accumulation of ROS in the mutant strain has a detrimental effect on its survival rate when exposed to an external oxidative stress.

## Discussion

In this study, our investigations unveiled that *melH* deletion disrupts coordination between ROS detoxification and thiol homeostasis. Our collective observations suggest an intricate narrative: *melH* deletion leads to heightened epoxide and aldehyde levels and accumulation of purine and quinolinic acid metabolites. Subsequently, increased levels of WhiB3 sensitize mycobacteria to oxidative stress through reduction in cellular levels of thiol buffers, EGT and MSH. Inability to clear ROS further disrupts the delicate balance of cellular redox homeostasis, overwhelming the antioxidant defense mechanisms of the mutant (Figure 8).

**Figure 8.**
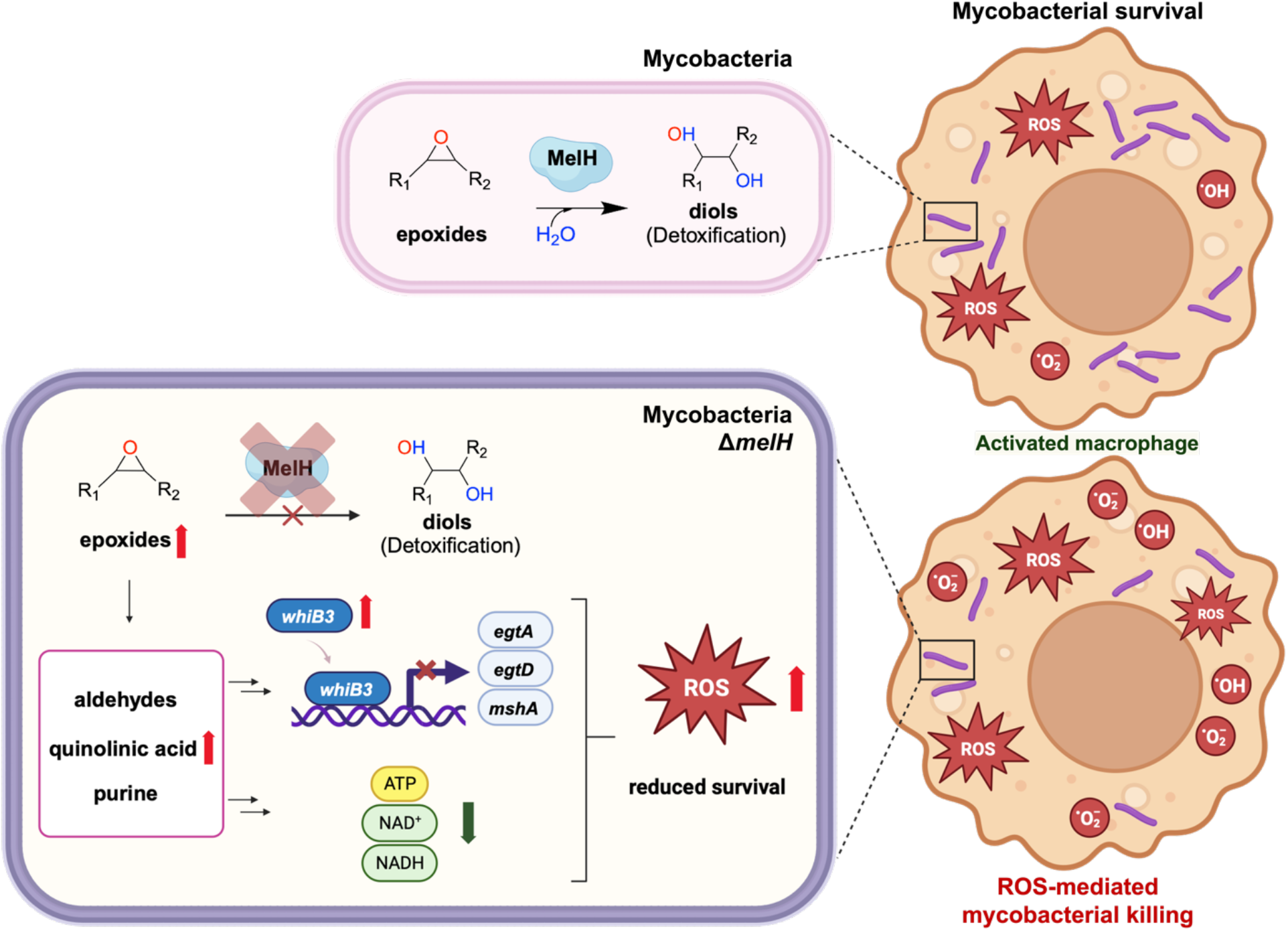
Model for metabolic consequences of *melH* deficiency. In *Mtb* and *Mm*, *melH* encodes MelH (epoxide hydrolase b), which catalyzes hydrolysis of epoxides and aids mycobacterial survival within host macrophages. In the absence of functional MelH in the Δ*melH* mutant, there is an accumulation of epoxides or their precursors, leading to the buildup of aldehydes. The deletion of *melH* results in elevated expression of *whiB3* and decreased production of the redox buffer EGT. In addition, deletion of *melH* causes a decrease in intracellular levels of NAD^+^, NADH, and ATP. All of these changes affect bioenergetic homeostasis and bacterial growth and ultimately sensitize the mutant to oxidative stress.

The natural substrate for MelH remains to be identified. The structure of *Mtb* MelH has been solved, revealing a relatively small and hydrophobic active site in a classic αβ hydrolase fold and substrate selectivity is altered compared to mammalian epoxide hydrolases (6, 8). Our metabolomics analysis did not reveal an accumulated epoxide substrate, therefore we investigated several epoxides as possible substrates with varying aromatic moieties and found that MelH efficiently catalyzes hydrolysis of styrene oxide and vitamin K1 2,3-epoxide into the corresponding diol, and does not catalyze hydrolysis of epoxides containing a biphenyl or two aromatic substituents (Table 2). Considering all the data in combination, the natural substrates of MelH are most likely aromatic species generated through metabolite breakdown and can be hindered epoxides like Vitamin K.

It is important to note that both epoxides and aldehydes can be volatile, are reactive electrophiles, and in the case of aldehydes, exist in hydrated forms. These factors may explain why we did not observe directly the accumulation of epoxides or aldehyde species other than *p*HBA by mass spectrometry. We did observe related metabolites, for example, an increased abundance of quinolinic acid (QA) in Δ*melH Mm* (Figure S8C). QA is a product of 2-amino-3-carboxymuconate-semialdehyde, an unstable compound that can be non-enzymatically transformed to QA (40). Hence, it is possible that 2-amino-3-carboxymuconate-semialdehyde is also one of the accumulated aldehydes in Δ*melH Mm*.

In addition, we found an accumulation of other metabolites in the cytokinin pathway, specifically N-isopentenyladenine and adenine in Δ*melH Mm*. This suggests that *melH* might be involved in cytokinin metabolism in *Mtb*. While cytokinins are well-established adenine-based signaling molecules in plants, their function in *Mtb* remains unknown (41).

Darwin and coworkers investigated the effects of accumulation of *p*HBA, a major product of cytokinin breakdown, on gene expression in mycobacteria (29–31, 41). In their RNA-Seq comparisons of WT *Mtb* treated with *p*HBA versus untreated WT, *p*HBA exhibited no influence on the expression of *whiB3*, *egtA*, *egtD*, and *mshA* genes. Thus formation of *p*HBA is not directly responsible for *whiB3* upregulation. The inducer of *whiB3* upregulation may be an unidentified epoxide, or another accumulating species that acts through one of the many known regulators of *whiB3*, e.g., PhoPR, RegX3, or GlnR (42, 43).

Additional contributors to the *melH* oxidative stress phenotype may be cross-talk with sigma factors (SigH and SigE), two-component systems (SenX-RegX and DosR/S/T), or serine-threonine kinases (PknG), all of which are reported to be key mediators of the oxidative stress response in *Mtb* (44). Another possibility is that the accumulation of QA and isonicotinic acid contributes to the growth and oxidative stress-resistant phenotype we observed in the Δ*melH* mutant. Previous studies have shown that QA can form complexes with Fe (II), inducing ROS formation, especially hydroxyl radical (•OH), which, in turn, is responsible for DNA chain breakdown and lipid peroxidation (45).

Furthermore, QA is a product of tryptophan degradation, which is crucial to produce NAD^+^ (46). Thus there is a potential connection between *melH* deletion and perturbations in tryptophan metabolism. The perturbations in tryptophan metabolites correlate with the reduced levels of NAD+ and NADH observed upon *melH* deletion. Reduction of NAD+/NADH levels can still impact metabolic flux through redox reactions, despite maintenance of the NAD+/NADH ratio. A recent study by the Schnappinger lab determined that bacteriostatic levels of NAD+ depletion cause compensatory remodeling of NAD-dependent metabolic pathways without impacting NADH/NAD+ ratios (23). On the other hand, bactericidal levels of NAD+ depletion led to a disruption of NADH/NAD+ ratios, inhibition of oxygen respiration. All of which may lead to decreased ATP levels.

Our CFU results indicated that the changes in NAD levels occuring upon *melH* deletion are not sufficient to elicit alterations in bacterial survival. However, under oxidative stress conditions, these changes are adequate to induce shifts in bacterial physiology, resulting in reduced survival. These data point toward a role for *melH* deletion causing NAD(H) depletion, further contributing to an inability to maintain redox homeostasis.

In summary, the findings of this study provide insight into the role of *melH* and its potential interactions with other pathways in mycobacterial metabolism and highlight the importance of further investigations of MelH function for understanding mycobacterial intracellular survival.

## Materials and Methods

### Bacterial strains and culture conditions

The *melF:Tn* (NR-13643), *melG:Tn* (NR-13645), and *melH:Tn* (NR-18002) *Mtb* CDC1551 strains were obtained from BEI Resources (www.beiresources.org); and the ymm1 *Mm* strain was kindly provided by Dr. Jeffrey D. Cirillo. *Mtb* and *Mm* strains were grown at 37°C and 30 °C, respectively, in Middlebrook 7H9 medium (Difco) supplemented with 0.2% glycerol or 0.1 mM cholesterol, 0.5% bovine serum albumin, 0.08% NaCl, and 0.05% (volume/volume) tyloxapol (Sigma-Aldrich) and filter-sterilization. Alternatively, cultures were grown on Middlebrook 7H11 agar supplemented with 10% oleate-albumin-dextrose-NaCl (OADC) and 0.5% glycerol. If necessary, cultures were supplemented with sodium acetate, succinic acid, fumaric acid, butyric acid, malic acid, or palmitic acid to a final concentration of 50 μM as sole carbon source. The *melF:Tn*, *melG:Tn*, and *melH:Tn* CDC1551 strains were grown in the presence of 30 μg/mL kanamycin, whereas the Δ*melH Mtb* CDC1551 and Δ*melH Mm* ymm1 strains were grown in the presence of 50 μg/mL hygromycin B. All complemented strains were grown in the presence of 50 μg/mL hygromycin and 50 μg/mL kanamycin. For determination of the effect of the carbon source on *melH* function, *Mtb* and *Mm* were cultured in Middlebrook 7H9 medium supplemented with 10% OADC, 0.05% tyloxapol, and one of the following carbon sources: glycerol, sodium propionate, sodium pyruvate, cholesterol, oleic acid, sodium acetate, succinic acid, fumaric acid, butyric acid, malic acid, or palmitic acid to a final concentration of 50 μM.

### Cloning of Δ*melH* and Δ*melH* complemented strains of *Mtb*

Δ*melH Mtb* was generated through recombinase-mediated recombination of a hygromycin-resistance cassette PCR product flanked by sequences homologous to the ∼545-bp regions upstream and downstream of the *melH* (*Rv1938*) gene. *Mtb* CDC1551 cells were transformed with an episomal plasmid containing the RecET recombinase (pNit-recET-sacB-kanR) and were induced to express the recombinase by addition of isovaleronitrile (final concentration, 1 μM) for 24 h followed by addition of glycine (final concentration, 0.2 M) for 16 h. The electrocompetent cells prepared from the induced culture were transformed with 1 μg of PCR product and were recovered for 24 h in 7H9 medium supplemented with 1 mM L-arginine. The transformants were selected on Middlebrook 7H10 agar containing 0.5% glycerol, 10% OADC, 50 μg/mL hygromycin, and 1 mM L-arginine and were validated by PCR. The validated strains were streaked on 7H11 agar supplemented with 0.5% glycerol, 10% OADC, 1 mM L-arginine, and 8.5% sucrose to select for clones that had lost the pNit-recET-sacB-kanR plasmid. The sucrose-resistant colonies were patched first onto 7H11 agar supplemented with 0.5% glycerol, 10% OADC, 1 mM L-arginine, and 30 μg/mL kanamycin and then onto the same agar without kanamycin. The kanamycin-sensitive strains were grown in 7H9 medium supplemented with 50 μg/mL hygromycin and 1 mM L-arginine to obtain the final Δ*melH Mtb* strain. For complementation of Δ*melH* strain, the target gene with a native promoter corresponding to 200-bp upstream of the first gene in the corresponding putative operon was cloned into a pMV306 integrating plasmid. The resulting construct was electroporated into the Δ*melH* mutant, and kanamycin-resistant transformants were selected.

### Cloning of Δ*melH* and Δ*melH* complemented strains in *Mm*

The predicted *melH* (MMAR_2866) gene and its flanking genomic regions, as annotated in the MycoBrowser portal, were amplified by PCR using *Mm* genomic DNA. The resulting upstream and downstream amplicons were fused by PCR with the primers listed in Table S1 and the fused PCR product was purified and digested with the *Hind*III restriction enzyme (New England BioLabs, Ipswich, MA) according to the manufacturer’s instructions. The *Hind*III-linearized p1NIL plasmid (Addgene plasmid number 20187) was ligated with the *Hind*III-digested PCR product, and the ligation mixture was introduced into *Escherichia coli* DH5α. Colonies bearing the p1NIL plasmid with the *melH* flanking regions from the *Mm* genome were selected on LB agar supplemented with kanamycin, and the sequence of the resulting p1NIL-*melH* plasmid was confirmed by DNA sequencing analysis. To construct the final suicide vector for allelic exchange, we introduced the p1NIL-*melH* plasmid with the pGOAL19 cassette (Addgene plasmid number 20190) using the protocol described by Parish and Stoker (47). The presence of the pGOAL19 cassette in the p1NIL-*melH* construct was confirmed by restriction digestion. The constructed plasmid was transformed into WT *Mm* by electroporation at 2500 mV, 1000 Ω, and 25 μF; and the *Mm* deletion mutant was selected after two rounds of homologous recombination, as previously described (47). Unmarked deletion mutants with in-frame deletions were obtained after two rounds of homologous recombination. Each transformed *Mm* mutant was cultured on 7H11 supplemented with 50 mg/L hygromycin for 2 weeks, and the single colony was then subcultured on 7H11 with 100 mg/L X-gal and 50 mg/L hygromycin for 1 week to select the single cross-over transformants (blue colonies). The blue colonies were selected and cultured on 7H11 supplemented with 2% sucrose and 100 mg/L X-gal for 1 week to obtain the deletion mutant.

### Central metabolite extraction

Mycobacteria were grown to an OD600 of 1.0. Approximately 1 × 10^8^ cells were collected by filtration on 0.22-μm nylon/polyvinylidene fluoride filters (Millipore GVWP02500) and transferred to 7H11 supplemented with 0.4% glycerol, sodium propionate, cholesterol, sodium pyruvate, or oleic acid as the sole carbon source. Plates were incubated for 6 days at 30 or 37 °C for bacterial replication and biomass production. The polar metabolites were extracted in prechilled (−40 °C) 2:2:1 methanol/acetonitrile/water, and cells were lysed six times by bead-beating for 30 s with incubation on ice for 30 s between pulses. Soluble extracts were filtered (Spin-X filter tubes) at 5000g for 5 min at 4 °C and then stored at −80 °C for liquid chromatography–mass spectrometry. The bacterial biomass of each sample was determined by measuring the residual protein content in the metabolite extracts.

### Metabolome analysis by liquid chromatography–mass spectrometry

The sample extract was fractionated by an Agilent 1290 Infinity II system equipped with 1) a Zorbax Eclipse Plus C18 column (1.8 μm; 2.1 by 50 mm) or 2) an InfinityLab Poroshell 120 HILIC-Z column (2.7 μM; 2.1 by 100 mm), or 3) a HiPlex H column (4.6 x 250mm, Agilent). For 1), reversed-phase separation was achieved using a gradient from solvent A (95% water, 5% acetonitrile, 0.1% formic acid) to solvent B (acetonitrile, 0.1% formic acid) as follows: 100% A held for 1.2 min, 100% B in 20.8 min, 100% B held for 1 min, 100% A in 0.1 min, and re-equilibration for 3 min (total run time, 26 min/sample). The flow rate was maintained at 200 μL/min for the duration of the run, the column was held at 35°C, and samples were held at 4°C. For 2) HILIC mode, separation was achieved for positive mode using a gradient from solvent A (10mM ammonium acetate in water) to solvent B (90% acetonitrile, 10% water, 10mM ammonium acetate) as follows: 98% B held for 0.8 min, 75% B in 19.2 min, 0% B in 3 min and held for 1 min, 98% B in 0.1 min, and re-equilibration for 3 min (total run time, 27 min/sample); For negative mode, separation was achieved using a gradient from solvent A (20 mM ammonium acetate in water, pH 9) to solvent B (acetonitrile) as follows: 100% B held for 0.8 min, 70% B in 17.2 min, 40% B in 4 min, 0% B in 1 min and held for 1 min, 100% B in 0.1 min, and re-equilibration for 3 min (total run time, 27 min/sample). The flow rate was maintained at 250 μL/min for the duration of the run, the column was held at 35°C, and samples were held at 4°C. For 3) HIPLEX mode, isocratic separation was achieved via flushing solvent at 200 μL/min (0.01% formic acid, 20% acetonitrile in water) for 20 minutes. The column was held at 50 °C. The column eluate was infused into a Bruker Impact II QTOF system with an electrospray ion source. Data were collected in both positive (A) and negative (B) ion mode, with the following settings: (A) Capillary voltage of 4,500 V; endplate offset of 500V; nebulizer gas pressure of 1.8 bar; dry gas flow rate of 8.0 L/min; dry temperature of 220 °C; MS spectra acquisition rate of 8 Hz; and m/z range of 50 to 1,500 Da). (B) Capillary voltage of 4,200 V; endplate offset of 500V; nebulizer gas pressure of 2.0 bar; dry gas flow rate of 8.0 L/min; dry temperature of 210 °C; spectra acquisition rate of 8 Hz; and m/z range of 50 to 1,500 Da).

### Analysis of metabolomics data

The data were analyzed using the Bruker Compass MetaboScape^®^ 2022b software (V9.0.1, Bruker Daltonics). Ion chromatograms were aligned, and high-resolution mass were re-calibrated using sodium formate clusters as reference mass. The metabolites were identified by comparing the accurate m/z (mass tolerance < 5 ppm) with libraries including HMDB library, Mtb LipidDB library and MycoMass library, and the MS/MS spectra with libraries including MetaboBASE Personal Library (Bruker), MSDIAL library, and HMDB library. Then the average peak intensities and the standard deviations from three biological replicates were calculated. Pathway map was plotted using *Mycobacterium tuberculosis* H37Rv metabolic map diagram (Biocyc.org). MetaboAnalyst (V5.0) was used to support metabolic pathway analysis (integrating pathway enrichment analysis and pathway topology analysis).

### Determination of *Mtb* and *Mm* susceptibility to oxidants

WT *Mtb* (CDC1551); *melF:Tn*, *melG:Tn*, *melH:Tn*, and Δ*melH Mtb*; WT ymm1 and Δ*melH Mm*; and their respective complemented strains were exposed to CHP or H_2_O_2_. The strains were grown to 0.6–0.8 OD_600_ in 7H9 medium supplemented with 0.4% glycerol or sodium propionate. For each strain, 2 × 10^6^ cells were transferred into each well of a 96-well plate. H_2_O_2_ (10 mM) or CHP (5 mM) was added to the wells, and the plates were incubated at 37°C for 30 mins. Control cells (1 × 10^6^) were not treated with either oxidant. The treated cells were subsequently plated on 7H11 agar plates supplemented with 10% OADC. Colonies were enumerated after 3 weeks of incubation at 37 °C. The survival of the mycobacterial strains was expressed as percentage survival relative to mycobacterial strains with no treatment.

### Measurement of endogenous ROS in *Mtb* and *Mm*

Exponentially growing WT *Mtb* (CDC1551); *melF:Tn*, *melG:Tn*, *melH:Tn*, and Δ*melH Mtb*; WT ymm1 and Δ*melH Mm;* and their respective complemented strains cultured in 7H9 medium supplemented with 0.4% glycerol or sodium propionate were treated with CellROX Green (Life Technologies; final concentration, 5 μM) for 30 min at 37 °C. The cells were pelleted, and the supernatant was discarded. The cells were washed with 7H9 medium to remove any extracellular CellROX Green. The washed cells were resuspended in medium and analyzed on a plate reader at excitation/emission wavelengths of 485/565 nm.

### Measurement of change in membrane potential

The changes in membrane potential of WT ymm1 *Mm*, Δ*melH Mm,* and the Δ*melH* complemented strain were determined using a BacLight Bacterial Membrane Potential Kit (ThermoFisher Scientific) according to the manufacturer’s instructions. In short, 1 mL aliquots of culture at an OD_600_ of 0.8 were treated with 3,3′-diethyloxacarbocyanine iodide (3 mM) and then CCCP (20 mM). DMSO-treated and -untreated bacilli were used as controls. Aliquots were incubated for 30 min before analysis on a plate reader.

### Measurement of total ATP content

Whole cell pellet samples were collected from *Mm* cell cultures, aliquots of the suspensions were removed from the samples, and the sample was mixed with 3 mL of boiling Tris-EDTA (100 mM Tris, 4 mM EDTA, pH 7.75). Then the bacterial cells were lysed for 2 min with glass beads, heated at 100 °C for 5 min, and cooled on ice. Cellular debris was removed by centrifugation. Supernatants were collected, an equal volume of luciferase reagent (ATP Bioluminescence Assay Kit HS II, Roche) was added to each supernatant, and luminescence was measured. ATP reaction mixing and ATP measurement were performed with an ATP Colorimetric/Fluorometric Assay Kit (BioVision Research Products, Milpitas, CA) according to the manufacturer’s protocol.

### NADH and NAD^+^ Determination

*Mm* Cells were rapidly harvested (two 2-mL samples) and resuspended in 0.2 M HCl (for NAD^+^ determination) or 0.2 M NaOH (for NADH determination). The tubes were centrifuged at 12,535 x*g* for 1 min. The supernatant was removed, and the pellets were suspended in 300 mL of 0.2 M HCl (for NAD^+^ extraction) or 0.2 M NaOH (for NADH extraction). The resulting suspensions were placed in a 50 °C water bath for 10 min and then on ice to cool them to 0 °C. The extracts were then neutralized with 300 mL of 0.1 M NaOH (for NAD^+^ extraction) or 300 mL of 0.1 M HCl (for NADH extraction) added dropwise while vortexing. Cellular debris was removed by centrifuging at 12,535 x*g* for 5 min. Supernatants were transferred to new tubes, and intracellular NADH and NAD^+^ concentrations were measured by means of a very sensitive cycling assay (48). The assay was performed with a reagent mixture consisting of equal volumes of 1.0 M bicine buffer (pH 8.0), absolute ethanol, 40 mM EDTA (pH 8.0), and 4.2 mM 3-[4,5-dimethylthiazol-2-yl]-2,5-diphenyltetrazolium bromide and twice the volume of 16.6 mM phenazine ethosulfate, which had previously been incubated for 10 min at 30 °C. The following volumes were added to 1 mL cuvettes: 50 µL of neutralized extract, 0.3 mL of water, and 0.6 mL of the above-described reagent mixture. The reaction was started by adding 50 µL of yeast Alcohol dehydrogenase II (500 or 100 U/mL in 0.1 M bicine buffer [pH 8.0]). The absorbance at 570 nm was recorded for 10 min at 30 °C.

### Quantitative real-time PCR

Total RNA was extracted from *Mm* cells using the TRIzol Reagent (Takara, Japan). cDNA was synthesized using a PrimeScript reverse transcriptase reagent kit (Takara, Japan). mRNA expression was examined by quantitative real-time PCR using iTaq Universal SYBR Green Supermix (Bio-Rad) and a LightCycler480 detection system (Roche, Mannheim, Germany). The relative expression of indicated genes was analyzed by the 2-ΔΔCt method. For comparisons between WT and Δ*melH Mm*, the induction ratio for each gene was normalized to *Mm* 16s rRNA expression. The primer sequences used for PCR are listed in Table #S1.

### Nitrite measurement

The *Mm* cells were treated with acidified NaNO_2_ and SNAP, and 30 min later, the Griess reagent system was used to determine levels of nitric oxide production. NO_2_^−^ was determined by using NaNO_2_ as a standard. Briefly, 50 μL of each experimental sample was added to wells, and then 50 μL of 1% sulfanilamide in 5% phosphoric acid was added to each experimental sample and to wells containing the dilution series for the nitrite standard reference curve. Then 50 of μL of 0.1% aqueous *N*-1-napthylethylenediamine dihydrochloride was added to each well. Absorbance was measured at 550 nm immediately after addition of the reagents to the samples. The cells were subsequently plated on 7H11 agar supplemented with 10% OADC. Colonies were enumerated after 3 weeks of incubation at 37 °C. The survival of mycobacterial strains was expressed as percentage survival relative to mycobacterial strains with no treatment.

### Aldehyde measurement

Aldehyde reaction mixing and Aldehyde concentration measurements were performed with a Fluorometric Aldehyde Assay Kit (MilliporeSigma, cat. no. MAK141), which uses a fluorogenic dye that reacts with aldehydes to generate a fluorometric product, according to the manufacturer’s protocol.

### MelH substrate Assays

150 ng of MelH recombinant proteins were used in each reaction. The reaction mixture contained 50 μM of vitamin K1 2,3-epoxide, 2-biphenylyl glycidyl ether, styrene oxide, 1,4-naphthoquinone 2,3-epoxide, or (E)-1,3-diphenyl-2,3-epoxypropan-1-one as the substrate. The reaction volume was adjusted with 50 mM Tris-HCl pH 7.5 to 250 μL, and the reactions were incubated at 37 ◦C for 60 min. Reactions were stopped by adding 500 μL of ethyl acetate and vortexing. The samples were briefly centrifuged, and the upper organic phase was transferred to a clean tube, dried under a stream of nitrogen, and resuspended in 20 μL of CHCl3:CH3OH (2:1). Half of the suspension was analyzed by thin-layer chromatography (TLC) in n-hexane:diethyl ether:formic acid (70:30:2). The remaining samples were dried under a stream of nitrogen, washed with CHCl3, resuspended in CHCl3, and analyzed by 1H-NMR spectroscopy. The initial velocity of epoxy fluor 7 substrate (Cayman, USA) turnover was measured in the presence of 25 μM of the indicated epoxide at pH 7.5, 37 °C for 10 minutes. The rate of turnover was then compared to the negative control without the addition of a second epoxide. 50 μM of epoxy fluor 7 was incubated with 150 ng of MelH in 50 mM Tris-HCl pH 7.5. The fluorescence intensity was monitored by a plate reader (Ex/Em: 330/465 nm).

## Statistics

Statistical computations were performed with GraphPad Prism (ver. 9.0). Pairwise comparisons were performed using Student’s *t* test for normally distributed data. Multiple comparisons were performed using the one-way analysis of variance module of GraphPad.

## Supplementary materials

The following material is available online. Supplemental Table, Figures and Methods, Spectra.

## Data availability

Metabolomics datasets from this study are deposited in the Global Natural Products Social Molecular Networking (GNPS; http://gnps. ucsd.edu) under reference MassIVE #MSV000092681. The data that support the findings of this study are available either within this article and its supplementary information files or upon reasonable request to the corresponding author.

## Author contributions

Yu-Ching Chen: Conceptualization, Methodology, Validation, Formal analysis, Investigation, Software, Data curation, Writing-Original Draft, Visualization. Xinxin Yang: Conceptualization, Methodology, Formal analysis, Supervision (Microbiology activities). Nan Wang: Methodology, Formal analysis, Software (Suggested and demoed the use of MetaboAnalyst), Supervision (Mass Spec activities). Nicole Sampson: Conceptualization, Resources, Writing-Review & Editing, Supervision, Funding acquisition.

## Acknowledgements

This work was supported by NIH grant R01AI134054. The metabolomic experiments were conducted at the Stony Brook CASDA Mass Spectrometry Center. The authors would like to thank Dr. Jeffrey D. Cirillo for generously providing the ymm1 *Mm* WT strain as a gift. The authors would like to thank Kun-Lin Hsieh for acquiring NMR spectra. The authors would like to thank Dr. Tianao Yuan for valuable discussions. Special thanks to Dr. Xuejun Peng for metabolomics advice. Shearson Editorial Services (Cornwall, NY, USA) provided English language editing of the text of this paper. Figure 8 was created with BioRender.com.

