## Supplemental Information for "Uncovering the Roles of *Mycobacterium tuberculosis melH* in Redox and Bioenergetic Homeostasis: Implications for Antitubercular Therapy"

**Running title: *melH* Deletion Causes Redox and Bioenergetic Imbalance**

Yu-Ching Chen,[a] Xinxin Yang,[b] Nan Wang,[b] Nicole S. Sampson[b,c,#]

[a]Program in Biochemistry and Structural Biology, Stony Brook University, Stony Brook, NY 11794-5215

[b]Department of Chemistry, Stony Brook University, Stony Brook, NY 11794-3400

[c]Department of Chemistry, University of Rochester, Rochester, NY 14627-0216

[#]Corresponding author:

### Keywords

*Mycobacterium tuberculosis*, redox homeostasis, ergothioneine, aldehyde, bioenergetic homeostasis

**Table S1. List of oligonucleotide sequences used as primers in this study**

| Primer name | Sequence 5'-3' | Description |
|---|---|---|
| **hygromycin cassette F** | ATATATCATATGATAACTTCGTATAATGTATGCTATACGAAGTTATGGATCATC | For hygromycin cassette cloning |
| **hygromycin cassette R** | ATATATAAGCTTATAACTTCGTATAGCATACATTATACGAAGTTATAAGCTGG | For hygromycin cassette cloning |
| ***Mtb _melH_*F** | ATAACATATGGTGTCGCAGGTCCATCG | For *Mtb melH* pretein expression |
| ***Mtb _melH_*R** | ATAAAAGCTTTCACGGCCGCAGC | For *Mtb melH* pretein expression |
| ***Mm_whiB3_*F** | GCAGTTGCAGGGCCTAT | For *Mm whiB3* gene expression |
| ***Mm _whiB3_*R** | TTGAGCAGCAGGTCACG | For *Mm whiB3* gene expression |
| ***Mm _egtA_*F** | CCGACCATGCGGATCAAT | For *Mm egtA* gene expression |
| ***Mm _egtA_*R** | GCTATCTGGACAGTCTTTCCG | For *Mm egtA* gene expression |
| ***Mm _egtD_*F** | CCGCCCAAATGGTTCTATGA | For *Mm edtD* gene expression |

| | | |
|---|---|---|
| ***Mm _egtD*_R** | TAGGCGTCCACGTTGAAATC | For *Mm egtD* gene expression |
| ***Mm _mshA*_F** | TTCGAGGGTCTGGACAAGTA | For *Mm mshA* gene expression |
| ***Mm _mshA*_R** | ACAGCCAATAGTGCGAGTG | For *Mm mshA* gene expression |

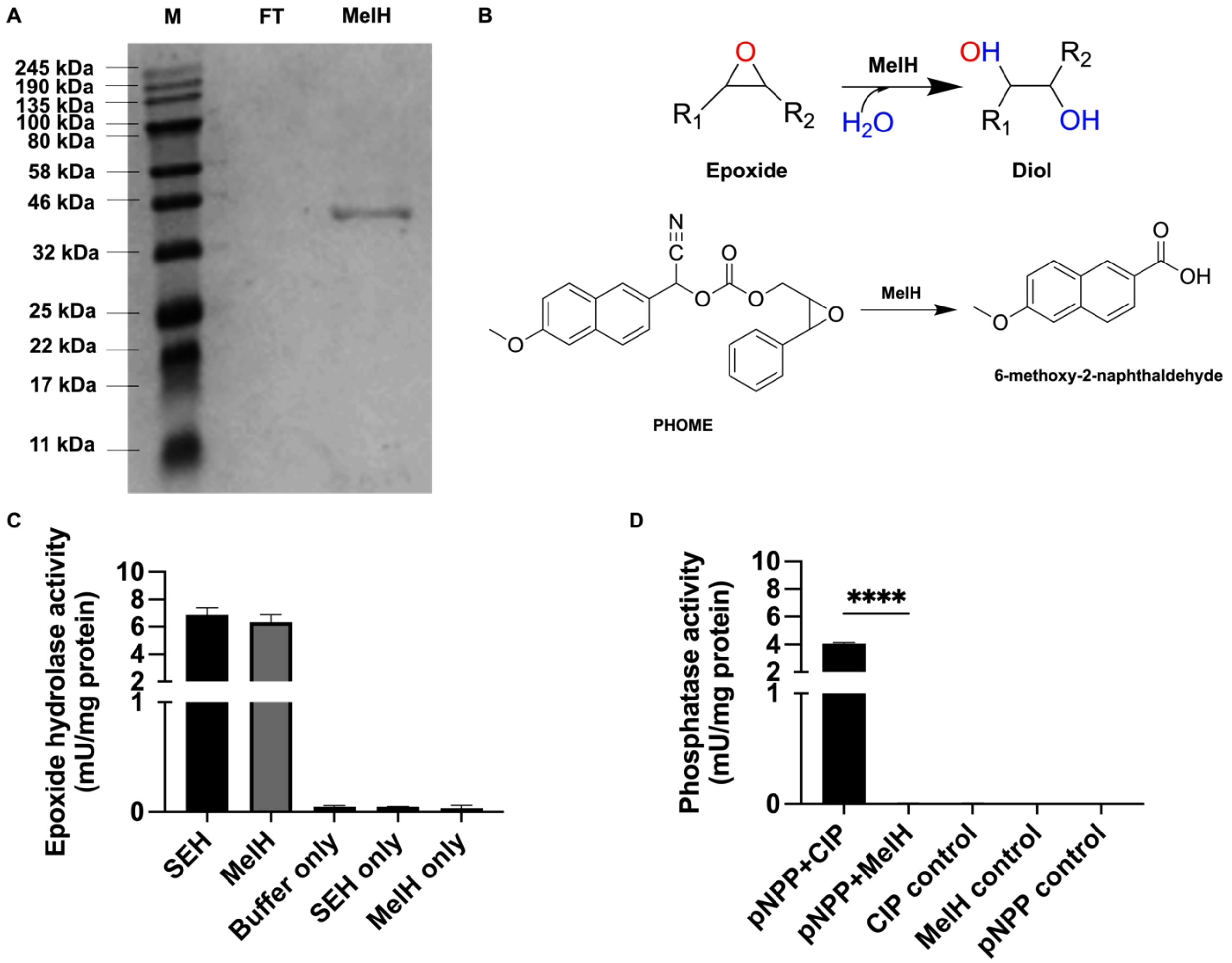


**Figure S1. Enzyme encoded by *melH* has epoxide hydrolase activity.**

(A) SDS-PAGE analysis of recombinant protein production in *Escherichia coli* BL21(DE3) pET28b-EphB$_{Mtb}$. A 6-His-tagged MelH (~40 kDa) recombinant protein was purified by Ni-nitrilotriacetic acid affinity and size-exclusion chromatography; and a protein marker (M), a flow-through control (FT), and MelH protein (MelH) were loaded onto the gel. (B) Reaction catalyzed by MelH and hydrolysis of artificial substrate PHOME, which was used for the epoxide hydrolase assay. (C) Epoxide hydrolase activity for sEH (positive control) and MelH (D) Analysis of alkaline phosphatase activity of calf intestinal alkaline phosphatase (CIP, control) and MelH. The substrate was *para*-nitrophenyl phosphate (pNPP), a generic phosphatase substrate whose dephosphorylated product, *para*-nitrophenol (pNP), is intensely yellow under alkaline conditions and absorbs at 405 nm. *P* values were determined by one-way analysis of variance using GraphPad Prism: ****$P < 0.0001$. Error bars indicate standard deviations of three replicate experiments.

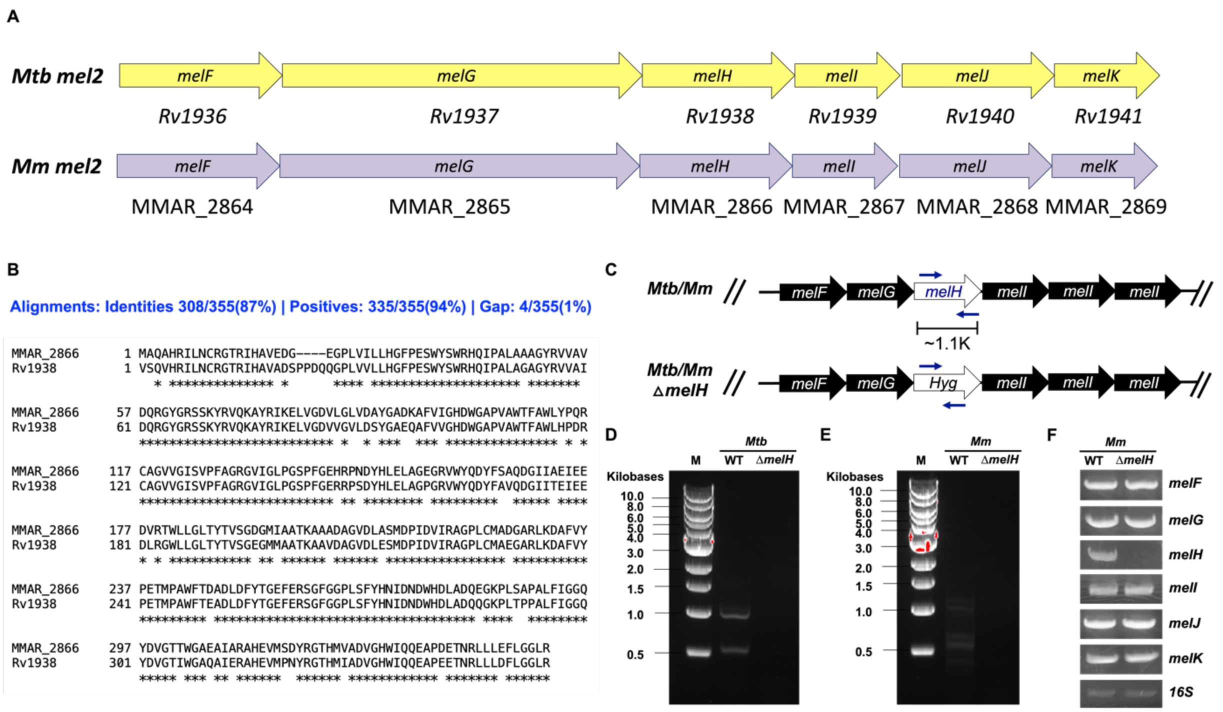


**Figure S2. Similarity of *Mtb melH* to *Mm melH*.**

(A) Schematic representation of the chromosomal region containing the *mel* genes of *mel2 loci* in *Mtb* and *Mm*. (B) The protein BLAST (BLASTp) sequence alignments of Mtb *Rv1938*-encoded MelH amino acid sequence with the amino acid sequence of Mm MMAR_2866-encoded MelH, revealed 87% identities, 94% positives, and 1% gap. (C) Schematic representation of *Mtb* and *Mm* genomic region containing *melH*, indicating the location of the primers used to validate allelic exchange and gene deletion of *melH*. (D) Validation of *Mtb melH* deleted mutant (Δ*melH* mutant) showing the PCR product amplimied by *melH* primers. (E) Validation of *Mm melH* deleted mutant (Δ*melH* mutant) showing the PCR product amplimied by *melH* primers. (F) Validation of *mel2* gene expressions in *Mm* Δ*melH* mutant by *melF, melG, melH, melI, melJ, melK* primers.

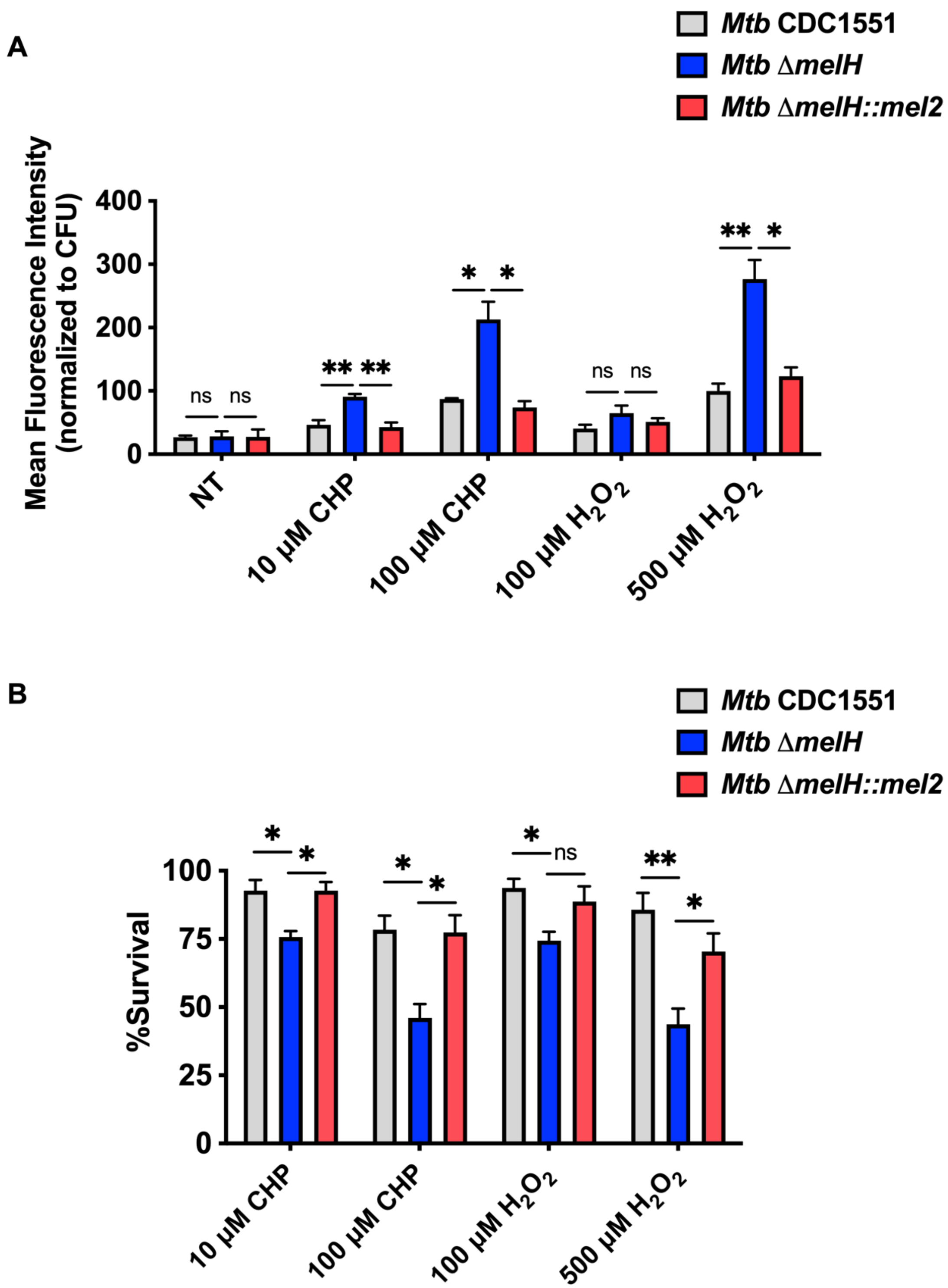


**Figure S3.** ***melH*** **deletion exacerbates effect of and increases susceptibility to oxidative stress.**

(A) Mean fluorescence intensity of CellROX Green in WT *Mtb* (CDC1551) the Δ*melH* mutant, and the Δ*melH* complemented strain (*melH*::*mel2*) in 7H9 medium containing glycerol as a single carbon source. (B) Percentage survival of *Mtb* determined by measuring bacterial CFU counts after a 48-h treatment with CHP or $H_2O_2$ in 7H9 medium containing glycerol as a single carbon source. *P* values were determined by one-way analysis of variance using GraphPad Prism: **P* $<$ 0.05, ***P* $<$ 0.01, ****P* $<$ 0.001; ns, not significant. Error bars indicate standard deviations of three replicate experiments.

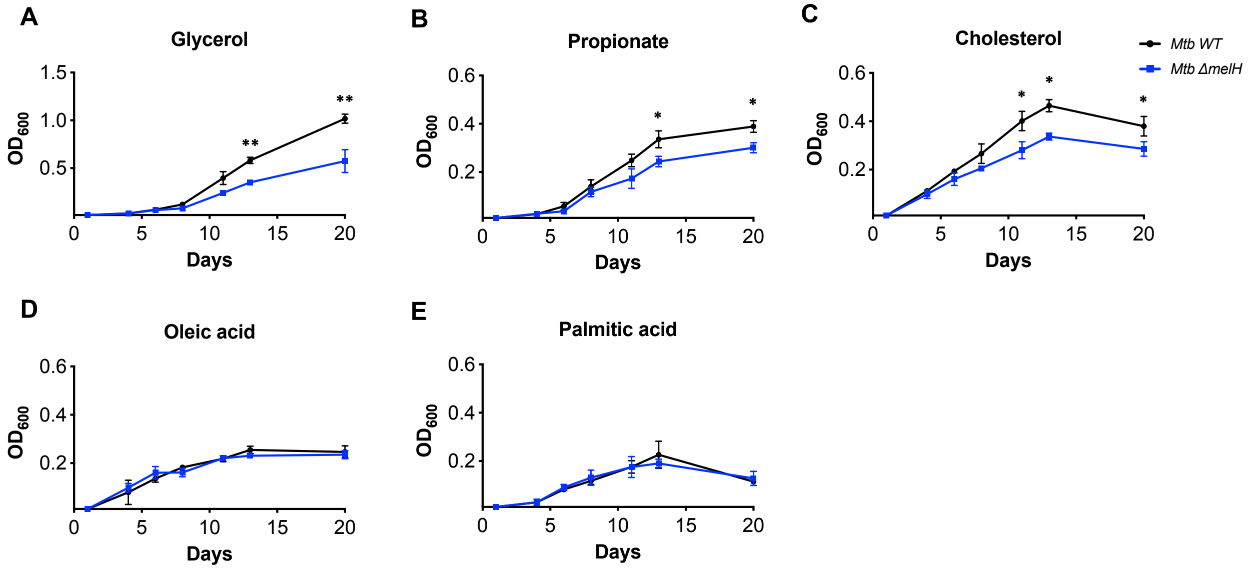


**Figure S4. Effect of *melH* deletion on *Mtb* growth rate depends on carbon source.**

Growth rates of WT *Mtb* (black lines) and the Δ*melH Mtb* mutant (blue lines) in minimal medium containing either (A) glycerol, (B) propionate, (C) cholesterol, (D) oleic acid, or (E) palmitic acid as a single carbon source. *P* values were determined by one-way analysis of variance using GraphPad Prism: *$P < 0.05$, **$P < 0.01$. Error bars indicate standard deviations. of three replicate experiments.

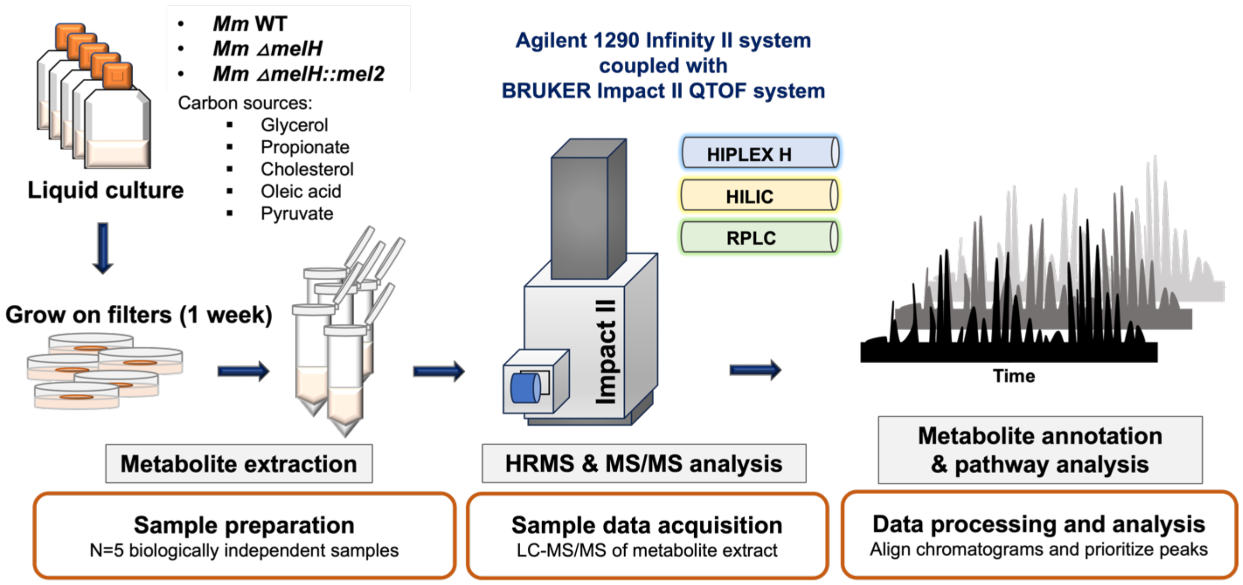


**Figure S5. Workflow and metabolite data analysis using untargeted metabolomic analysis.**

Overview of global, untargeted metabolomic workflow.

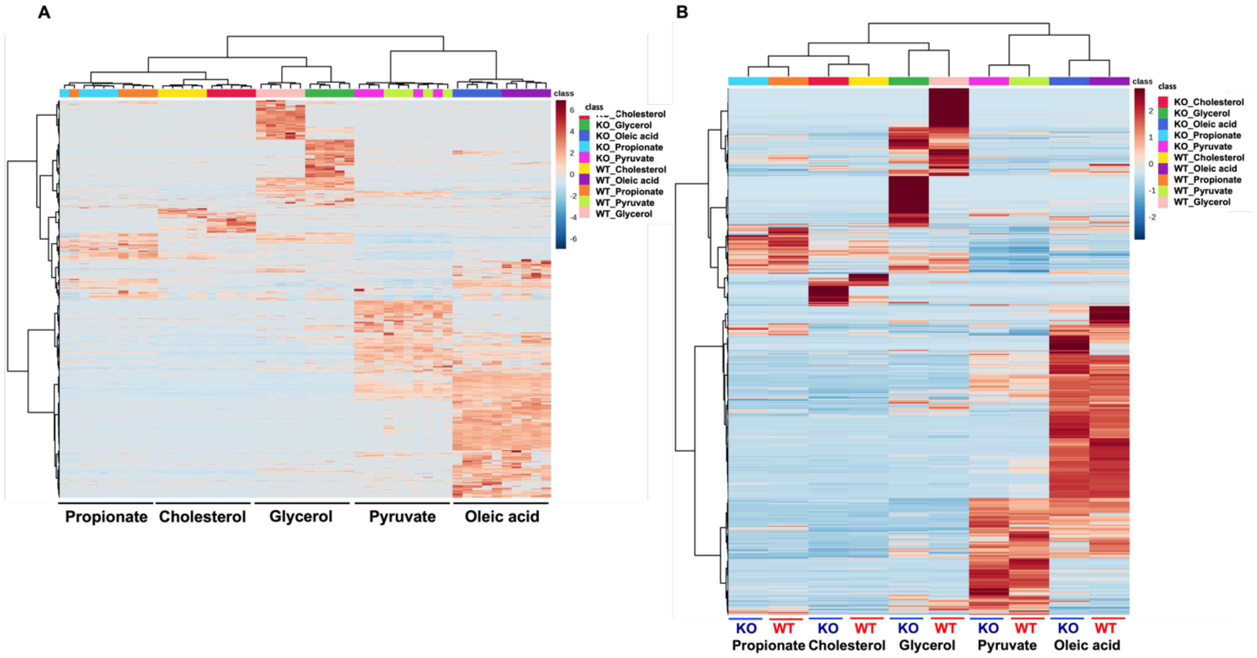


**Figure S6. Liquid chromatography–mass spectrometry analysis of putative metabolites in *Mm* demonstrates distinct metabolic differences in response to carbon catabolism.**

(A) Heat map showing ratios of metabolite concentrations in individual cell extracts prepared from Δ*melH Mm* (*N* = 5) versus concentrations in WT *Mm* (*N* = 5) cultured in the presence of glycerol, propionate, cholesterol, oleic acid, or pyruvate. (B) Heat map showing average ratios (*N* = 5) of metabolite concentrations in Δ*melH* versus concentrations in the WT grown on glycerol, propionate, cholesterol, oleic acid, or pyruvate.

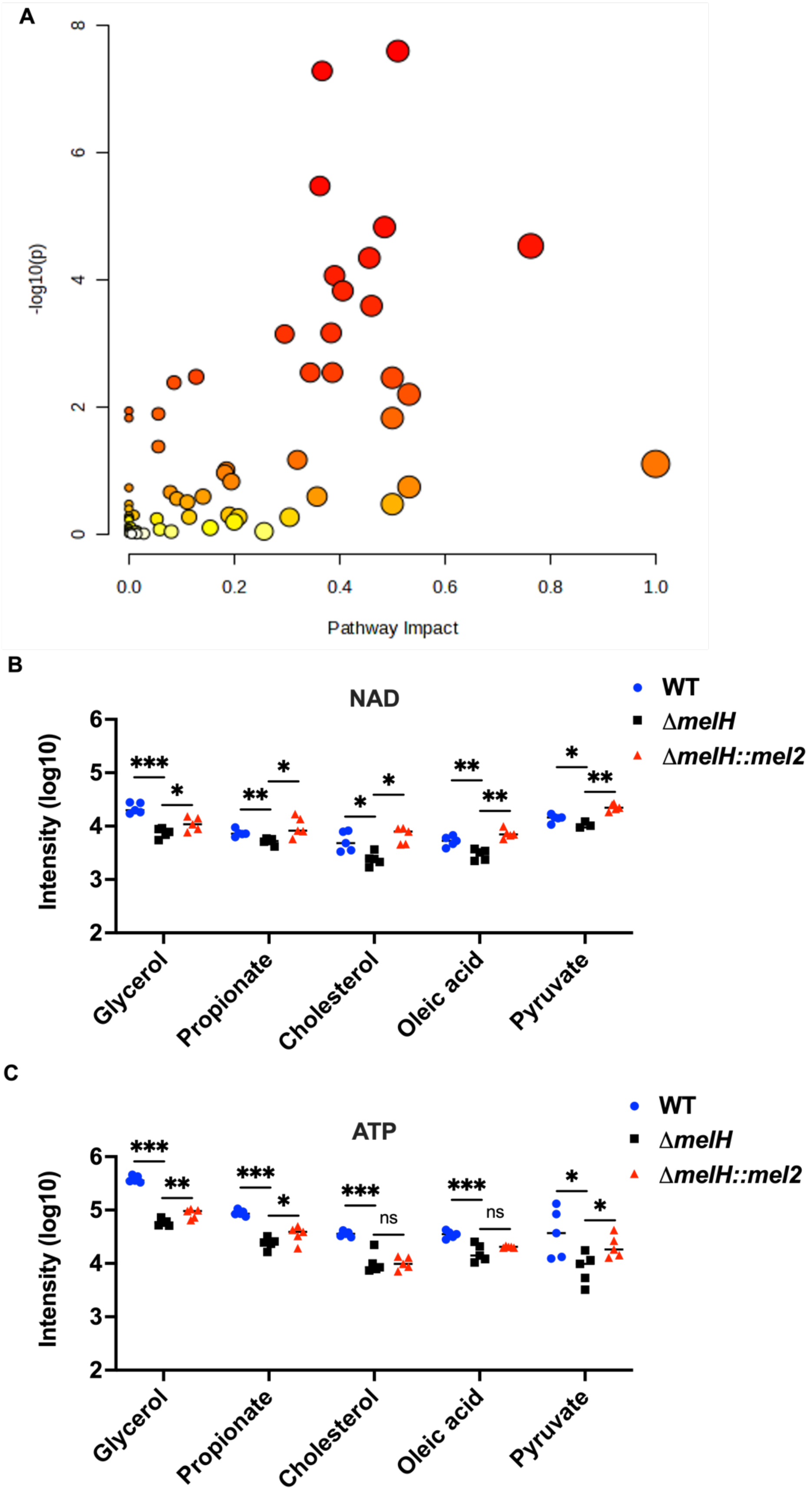
A
-log10(p)
Pathway Impact
B
NAD
Intensity (log10)
Glycerol
Propionate
Cholesterol
Oleic acid
Pyruvate
WT
ΔmelH
ΔmelH::mel2
C
ATP
Intensity (log10)
Glycerol
Propionate
Cholesterol
Oleic acid
Pyruvate
WT
ΔmelH
ΔmelH::mel2

**Figure S7. Metabolomic analysis of *melH*-deficiency *Mm.*** (A) Untargeted metabolite profiling assessed by MetPa enrichment analysis, based on changes in the relative abundance of metabolites (see also Table S1), indicate that melH has a profound effect on Mm metabolism. Corresponding levels of NAD (B), and ATP (C) in WT *Mm* ($N = 5$, blue dot), the Δ*melH Mm* mutant ($N = 5$, black square), and Δ*melH* complemented (*melH*::*mel2*) *Mm* ($N = 5$, red triangle). $P$ values were determined by one-way analysis of variance using GraphPad Prism: *$P < 0.05$, **$P < 0.01$, ***$P < 0.001$.

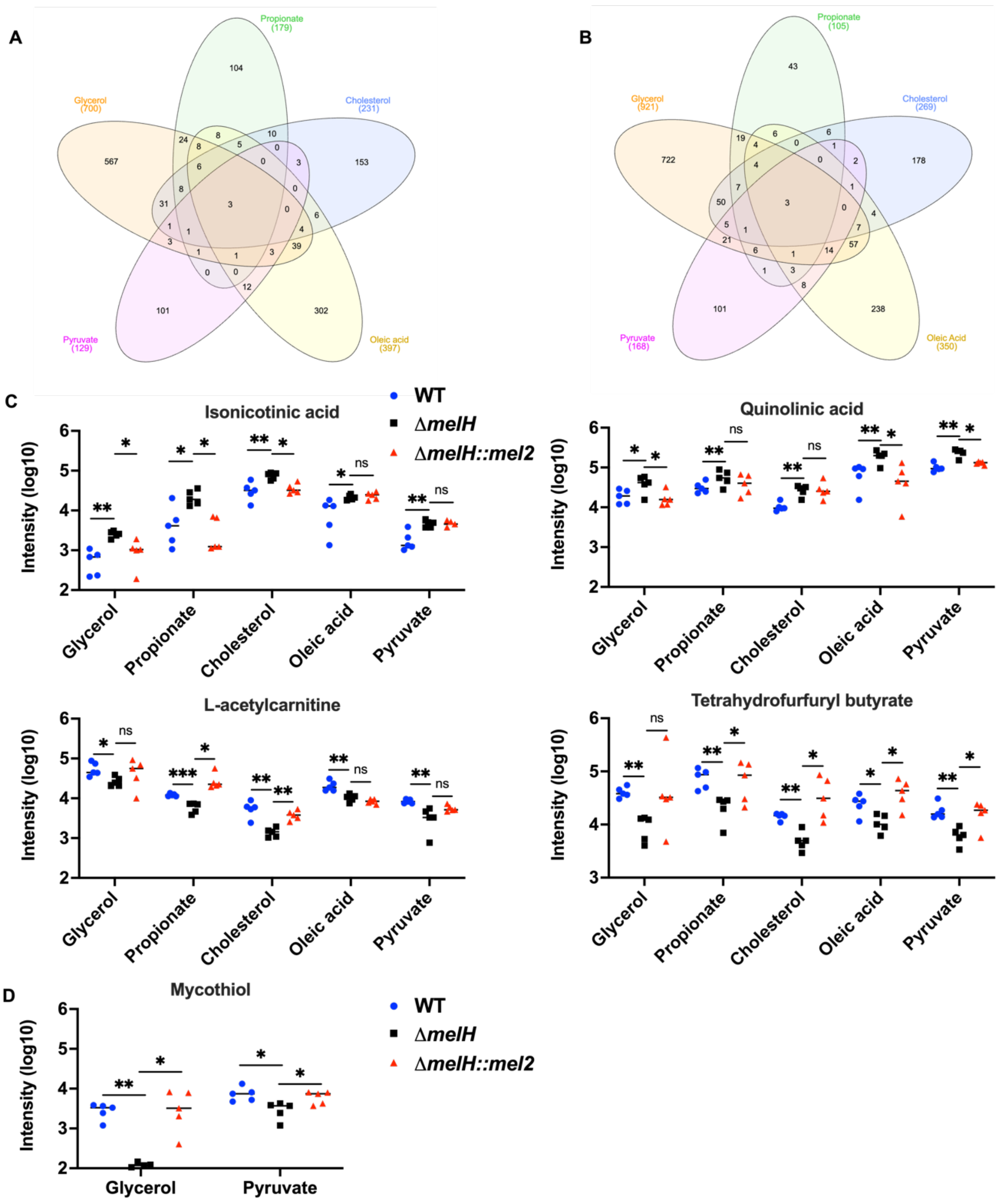
A
Propionate (179)
Glycerol (700)
Cholesterol (231)
Pyruvate (129)
Oleic acid (397)
B
Propionate (105)
Glycerol (921)
Cholesterol (269)
Pyruvate (168)
Oleic Acid (350)
C
Isonicotinic acid
Quinolinic acid
L-acetylcarnitine
Tetrahydrofurfuryl butyrate
Intensity (log10)
Glycerol
Propionate
Cholesterol
Oleic acid
Pyruvate
WT
ΔmelH
ΔmelH::mel2
D
Mycothiol
Intensity (log10)
Glycerol
Pyruvate
WT
ΔmelH
ΔmelH::mel2

**Figure S8. Metabolomic analysis of *melH*-deficiency *Mm*.**

(A) Venn diagram illustrating overlap of significantly increased metabolites (para-hydroxybenzaldehyde, Isonicotinic acid, Quinolinic acid) in Δ*melH Mm* grown with different carbon sources. (B) Venn diagram illustrating overlap of significantly decreased metabolites (L-acetylcarnitine, EGT, and tetrahydrofurfuryl butyrate) in Δ*melH Mm* grown with different carbon sources. (C) Corresponding levels of Isonicotinic acid, Quinolinic acid, L-acetylcarnitine, and tetrahydrofurfuryl butyrate in WT *Mm* ($N = 5$, blue dot), the Δ*melH Mm* mutant ($N = 5$, black square), and Δ*melH* complemented (*melH*::*mel2*) *Mm* ($N = 5$, red triangle). (D) Corresponding levels of MSH in WT *Mm* ($N = 5$, blue dot), the Δ*melH Mm* mutant ($N = 5$, black square), and Δ*melH* complemented (*melH*::*mel2*) *Mm* ($N = 5$, red triangle). *P* values were determined by one-way analysis of variance using GraphPad Prism: *$P < 0.05$, **$P < 0.01$, ***$P < 0.001$.

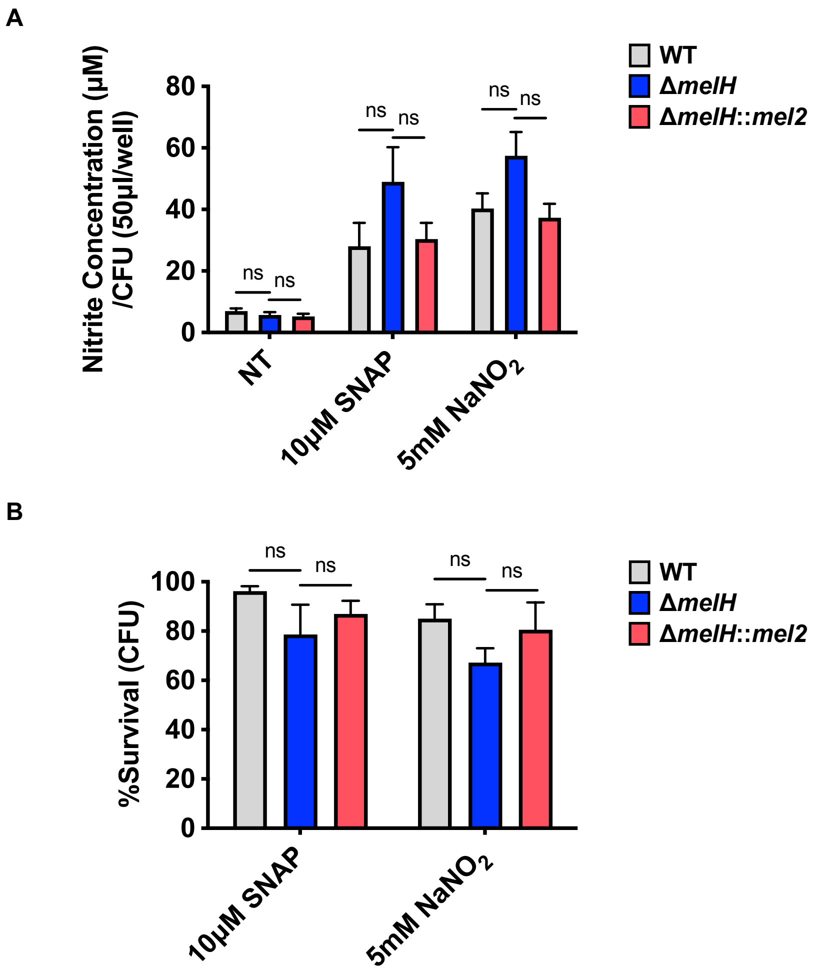


**Figure S9. Impact of *melH* deletion on nitrite production in *Mm*.**

(A) Nitrite production in WT, Δ*melH*, and Δ*melH* complemented (*melH*::*mel2*) strains of *Mm* grown in the presence of *S*-nitroso-*N*-acetylpenicillamine SNAP or $NaNO_2$ with propionate as the sole carbon source. The nitric oxide assay was conducted with the Griess reagent system. NT, no treatment. (B) Percentage survival of WT, Δ*melH*, and *melH*::*mel2 Mm* determined by measuring bacterial CFU counts after a 30-min treatment with SNAP or $NaNO_2$. *P* values were determined by one-way analysis of variance using GraphPad Prism; ns, not significant. Error bars indicate mean ± SD ($N = 3$ independent experiments).

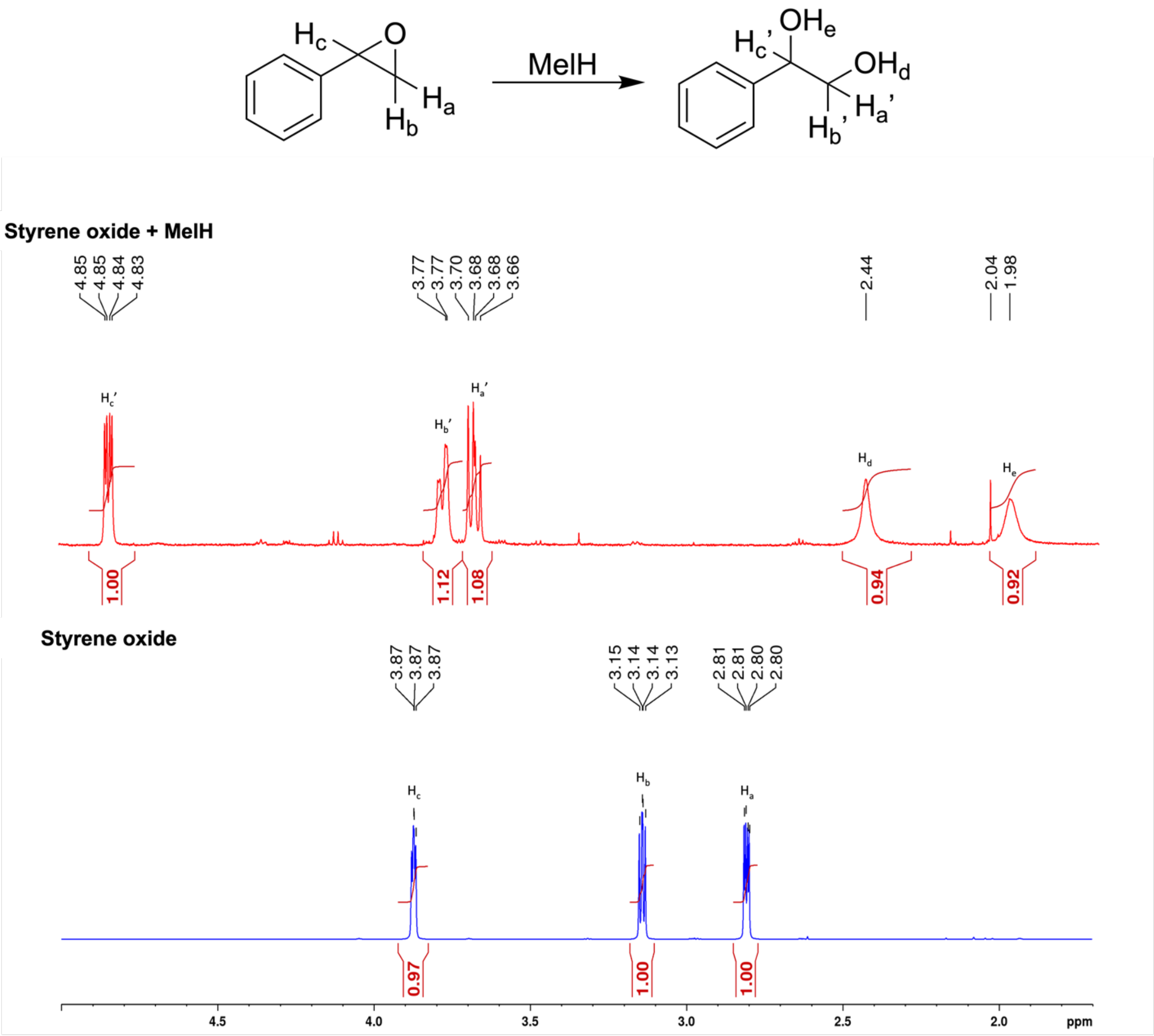


**Figure S10. MelH catalyzes hydrolysis of styrene oxide.**

$^{1}$H-NMR spectra of the substrate and product from the MelH catalyzed reaction with styrene oxide. Styrene oxide (50 µM) was incubated with MelH (150 ng) at pH 7.5, 37 °C for 1 hour. Top: Ethyl acetate extract of the reaction mixture containing the diol product. Bottom: Styrene oxide.

**Styrene oxide**

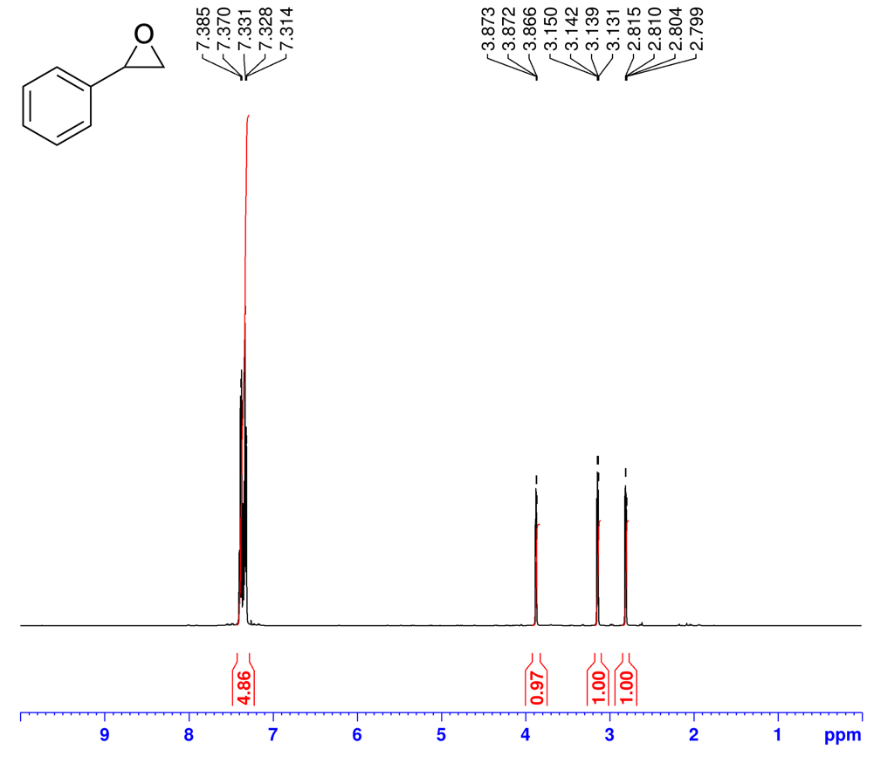


**Styrene oxide+MeIH**

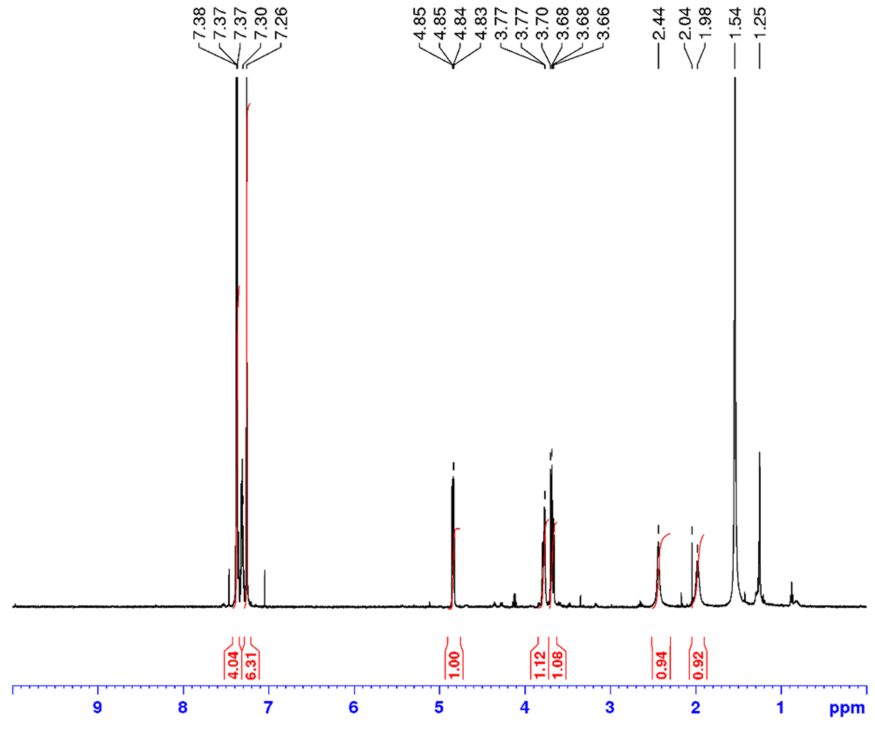

### 1,4-Naphthoquinone 2,3-epoxide

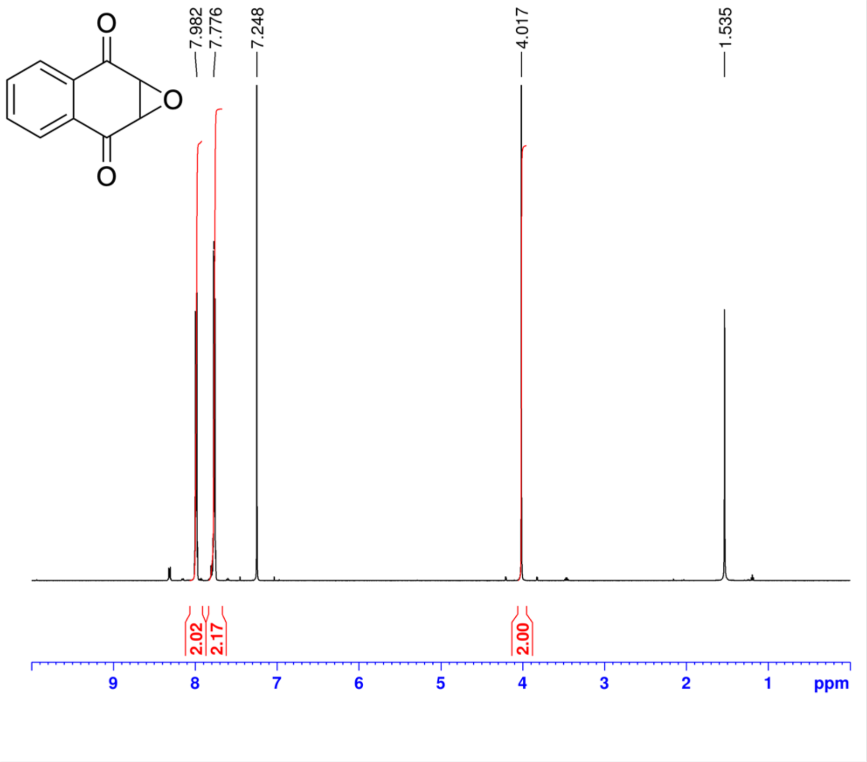


### 1,4-Naphthoquinone 2,3-epoxide+MeIH

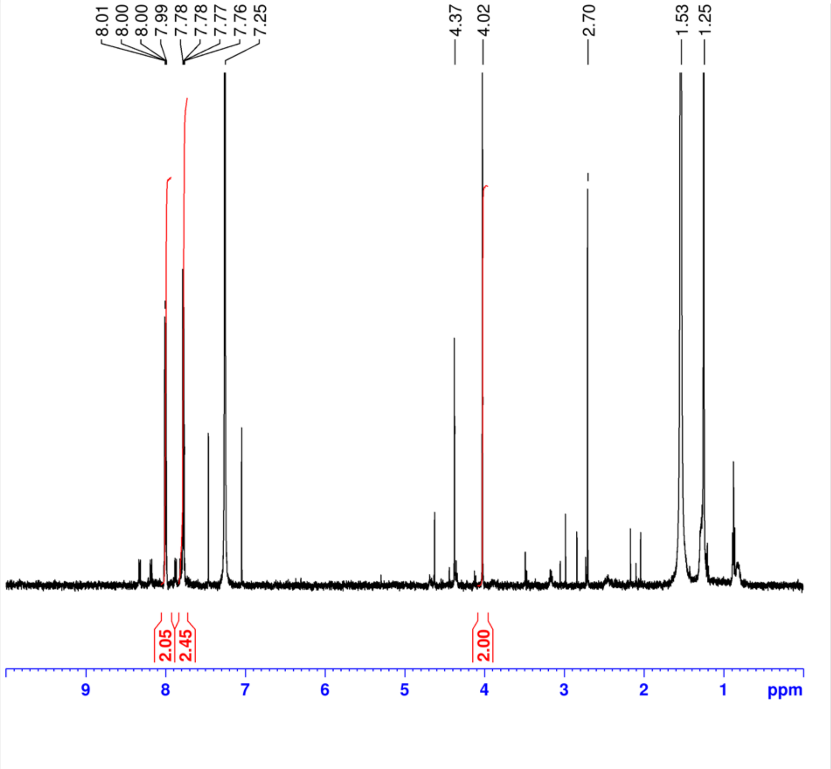

**(E)-1,3-Diphenyl-2,3-epoxypropan-1-one**

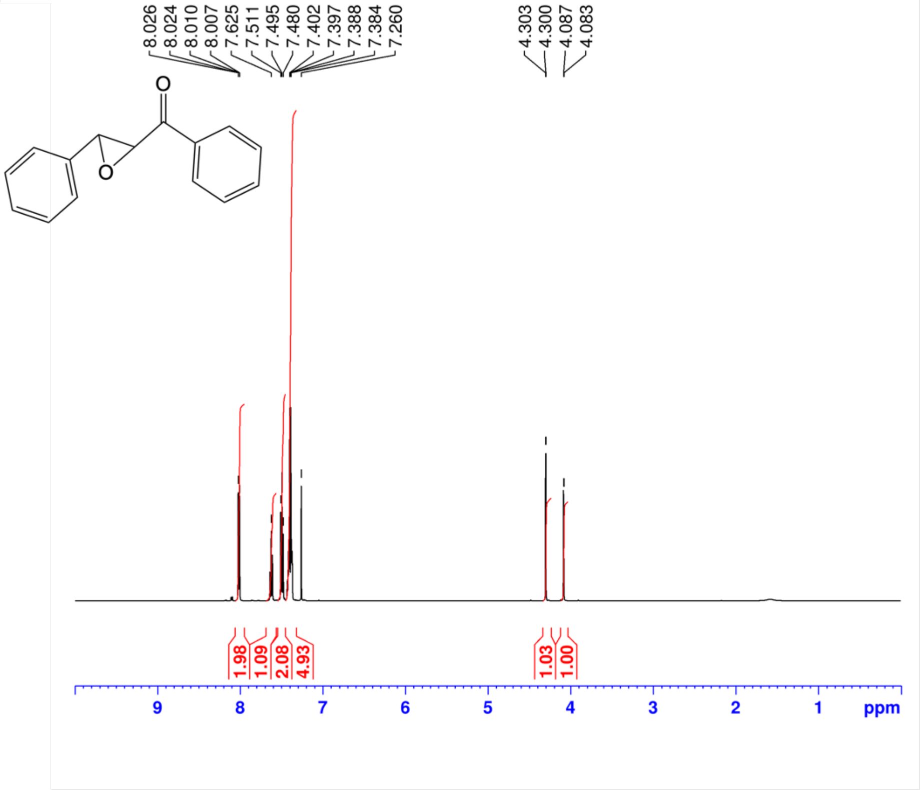


**(E)-1,3-Diphenyl-2,3-epoxypropan-1-one+MeIH**

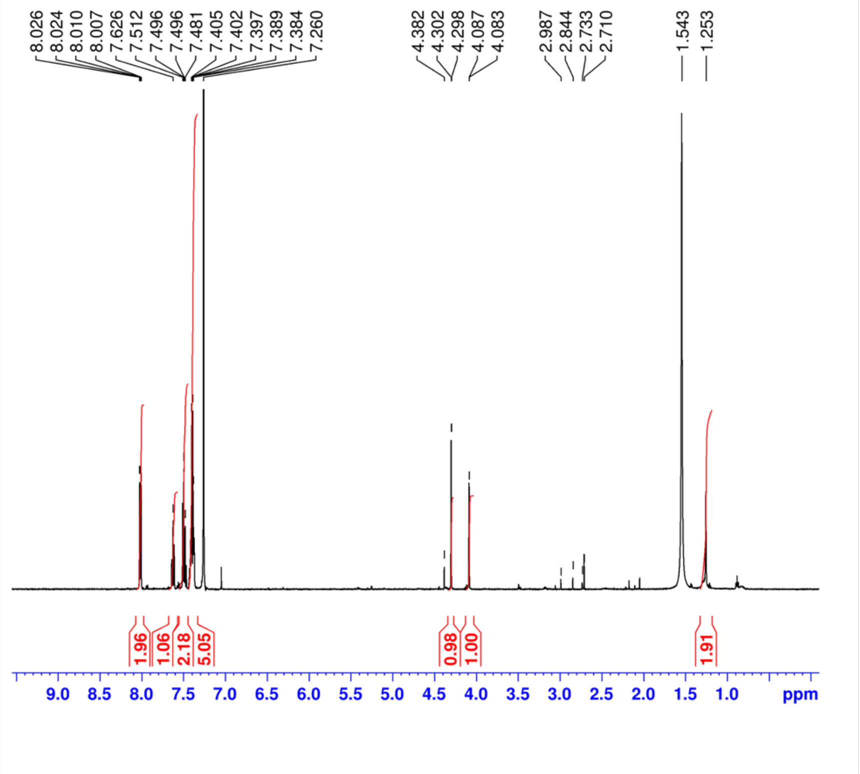

### 2-Biphenylyl glycidyl ether

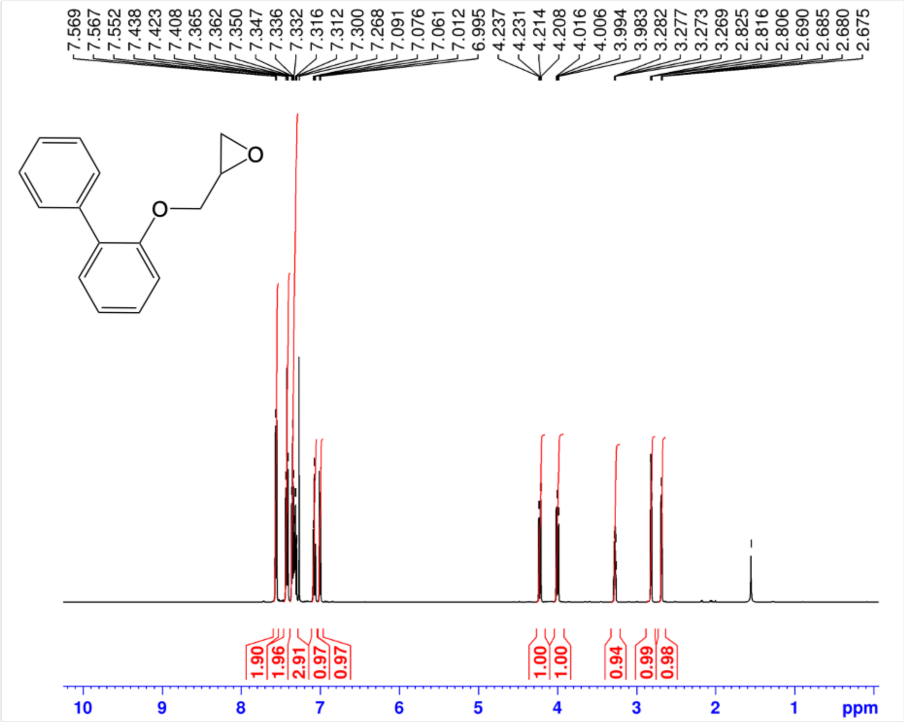


### 2-Biphenylyl glycidyl ether+MeIH

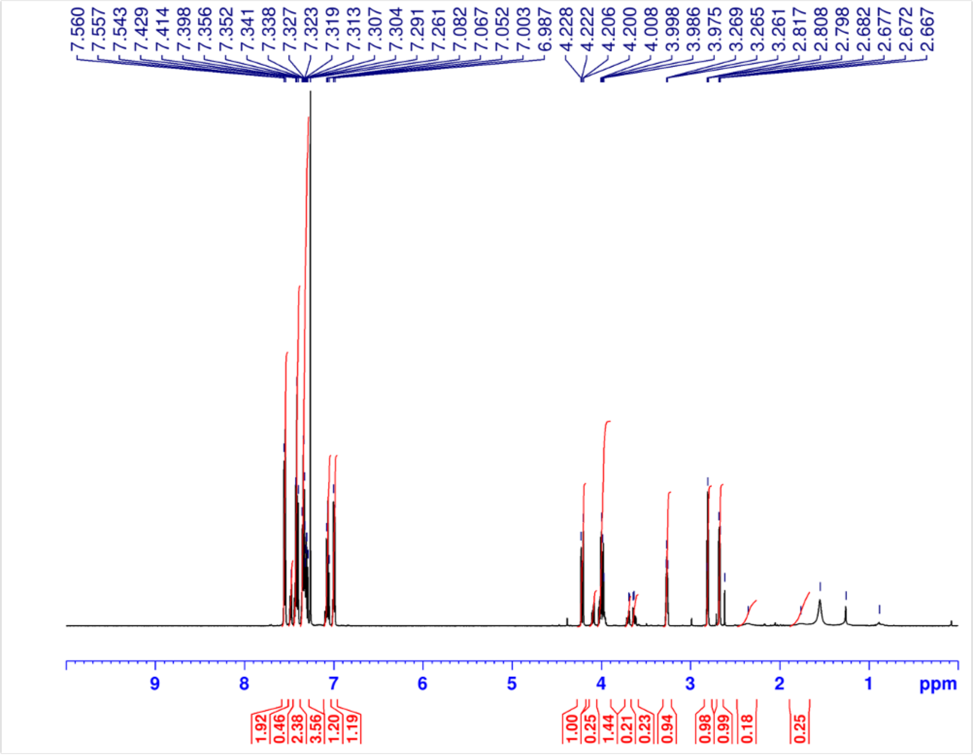

**Vitamin K**

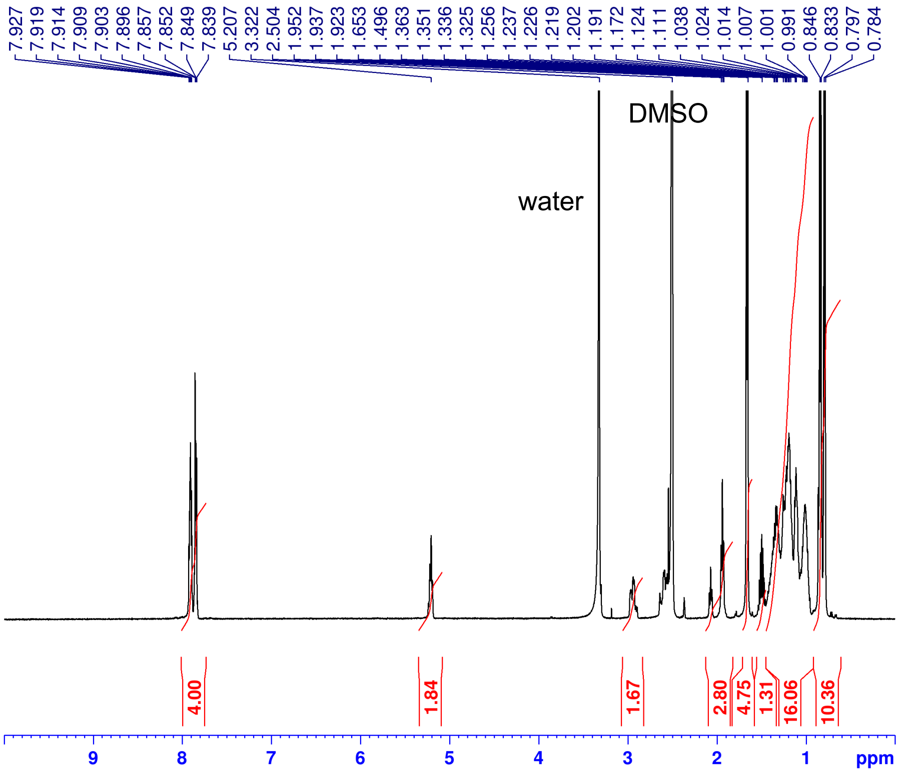


**Vitamin K + MelH**

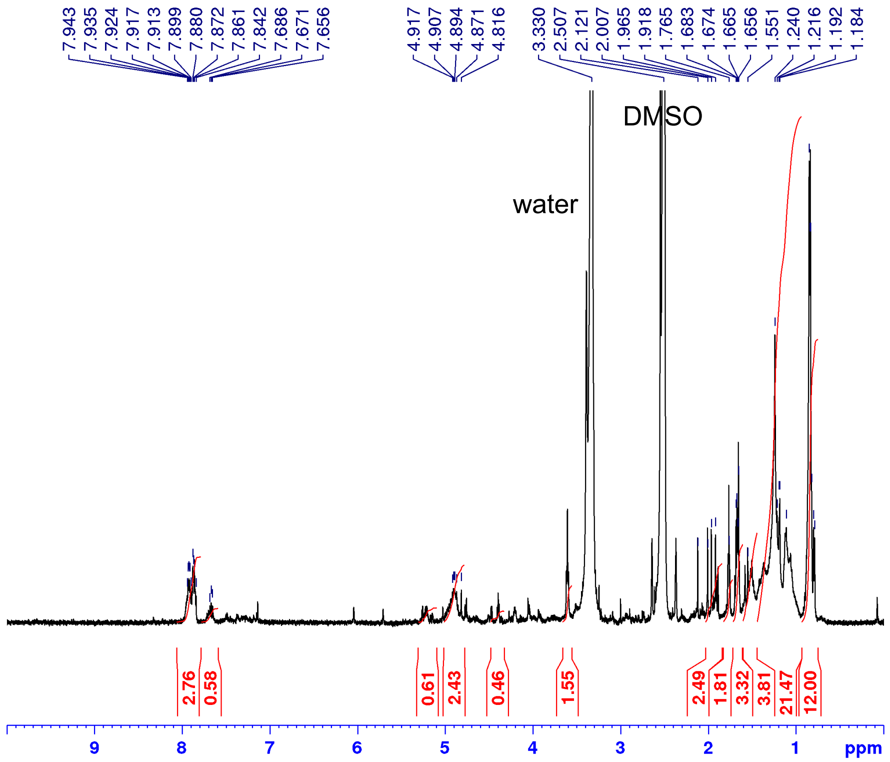
